# A targeted subgenomic approach for phylogenomics based on microfluidic PCR and high throughput sequencing

**DOI:** 10.1101/021246

**Authors:** Simon Uribe-Convers, Matthew L. Settles, David C. Tank

**Author notes:** Corresponding author (SUC). These authors contributed equally to this work (Simon Uribe-Convers, Matthew L. Settles).

## Abstract

Advances in high-throughput sequencing (HTS) have allowed researchers to obtain large amounts of biological sequence information at speeds and costs unimaginable only a decade ago. Phylogenetics, and the study of evolution in general, is quickly migrating towards using HTS to generate larger and more complex molecular datasets. In this paper, we present a method that utilizes microfluidic PCR and HTS to generate large amounts of sequence data suitable for phylogenetic analyses. The approach uses a Fluidigm microfluidic PCR array and two sets of PCR primers to simultaneously amplify 48 target regions across 48 samples, incorporating sample-specific barcodes and HTS adapters (2,304 unique amplicons per microfluidic array). The final product is a pooled set of amplicons ready to be sequenced, and thus, there is no need to construct separate, costly genomic libraries for each sample. Further, we present a bioinformatics pipeline to process the raw HTS reads to either generate consensus sequences (with or without ambiguities) for every locus in every sample or—more importantly—recover the separate alleles from heterozygous target regions in each sample. This is important because it adds allelic information that is well suited for coalescent-based phylogenetic analyses that are becoming very common in conservation and evolutionary biology. To test our subgenomic method and bioinformatics pipeline, we sequenced 576 samples across 96 target regions belonging to the South American clade of the genus *Bartsia* L. in the plant family Orobanchaceae. After sequencing cleanup and alignment, the experiment resulted in ∼25,300bp across 486 samples for a set of 48 primer pairs targeting the plastome, and ∼13,500bp for 363 samples for a set of primers targeting regions in the nuclear genome. Finally, we constructed a combined concatenated matrix from all 96 primer combinations, resulting in a combined aligned length of ∼40,500bp for 349 samples.

## Introduction

Advances in high-throughput sequencing (HTS) have allowed researchers to obtain large amounts of genomic information at speeds and costs unimaginable only a decade ago. The fields of phylogenetics and population genetics have benefitted greatly from these advancements, and large phylogenomic and population genomic datasets are becoming more common [1-3]. Driven by the need to generate homogenous, informative, and affordable multilocus datasets, we present a new approach for obtaining affordable large, multilocus datasets for phylogenetic and population genetic studies, based on microfluidic PCR amplification and HTS. Microfluidic PCR technology has been used extensively in the fields of cancer research [e.g., 4,5], genotyping of single nucleotide polymorphisms (SNP) [e.g., 6-8], gene expression [e.g., 9-11], and targeted resequencing [e.g., 12,13] but, to our knowledge, has not yet been used to assemble molecular phylogenetic datasets for systematic studies [but see 14 for a discussion of its potential use for phylogenetics]. This approach uses the Fluidigm Access Array System (Fluidigm, San Francisco, CA, USA) and two sets of PCR primers to simultaneously amplify 48 target regions across 48 samples, incorporating sample-specific barcodes and HTS adapters (2,304 amplicons per microfluidic array). This four-primer PCR approach circumvents the need to construct genomic libraries for every sample, avoiding the high costs and time requirements involved in library preparation. Furthermore, by using a dual barcoding strategy, we are able to multiplex on a single Illumina MiSeq run 24 microfluidic arrays, representing two distinct sets of 48 target regions (plastome and nuclear, in this experiment) across 576 samples (55,296 distinct amplicon sequences) from the South American clade of the plant genus *Bartsia* L. (Fig. 1) and its close relatives in Orobanchaceae, demonstrating the power of this approach for species-level phylogenetics.

**Fig.1.**
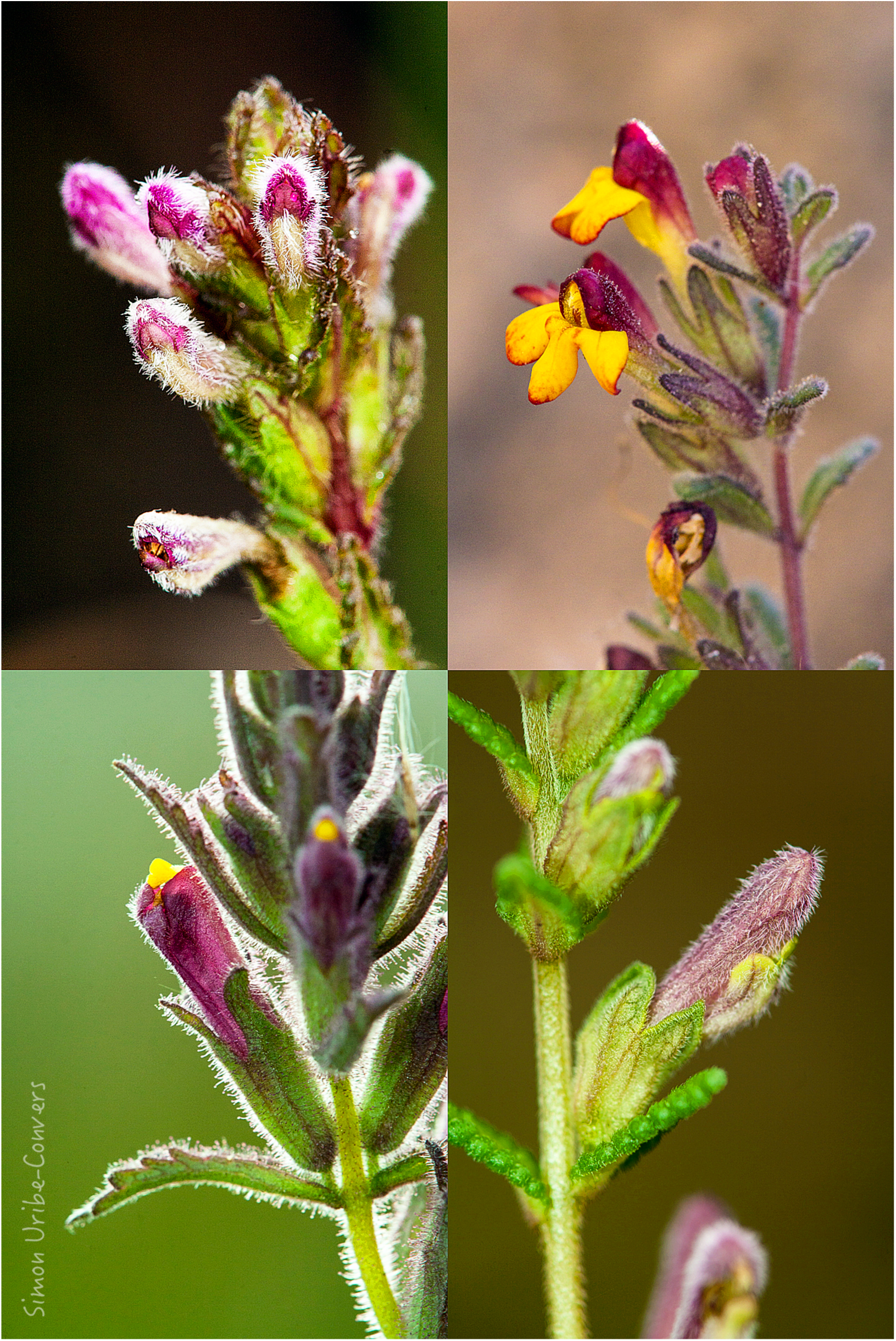
Floral diversity in the South American *Bartsia* clade.

The sections from top left to bottom right are: *Strictae, Diffusae, Laxae*, and *Orthocarpiflorae*.

In plant phylogenomics, there has been a special focus on the chloroplast genome, also know as the plastome, given its phylogenetic informativeness at all taxonomic scales [e.g., 15-18], the straightforwardly interpreted results due to its non-recombining nature, conserved gene order, and gene content [19], and its historical importance since the beginning of the field [e.g., 20]. Large datasets have been produced with approaches that have involved massively parallel sequencing [e.g., 17], a compilation of coding regions from both whole plastome sequences [e.g., 18] and targeted approaches [e.g., 21], transcriptomics [e.g., 22], RNA hybridization or capture probes [e.g., 3], and long range PCR coupled with HTS [e.g., 23]. Because the chloroplast genome evolves relatively slowly, ∼3–5 times slower than the nuclear genome in plants [24-26], the power of these datasets for phylogenetic studies lies in their size; at ∼150 kilobases (kb), plastome datasets can provide phylogenetic resolution from the interspecific level [e.g., 17,27] to the level of major clades [e.g., 18,28]. However, because it is inherited as a single unit, plastome sequences only provide information from a single locus, and although often well-supported, phylogenies based solely on plastome-scale datasets may be misleading because of the well known problem of gene tree-species tree discordance [29]. This may be especially problematic at low-taxonomic scales where processes such as coalescent stochasticity and gene flow may be more prevalent. Thus, data from multiple independently evolving loci is necessary to fully understand the evolutionary history of a group of organisms, and to take full advantage of the emerging species tree paradigm made possible by the integration of population genetic processes into phylogenetic reconstruction via the multispecies coalescent [30-32].

For the nuclear genome, phylogenomic datasets have been obtained in plant systems using genome skimming [e.g., 33], sequence capture [e.g., 34,35], and restriction-site associated DNA—RADseq [e.g., 36]. Likewise, the field of phylogenomics in animals has advanced with datasets obtained with targeted amplicon sequencing (TAS) in Pancrustacea [37] and North American tiger salamanders [38], GBS in butterflies [39], fish [40], and beetles [41]. However, genome-scale datasets for animal phylogenetics has been most heavily impacted by sequence capture approaches focused on ultraconserved genomic elements (UCEs) at various taxonomic scales, e.g., vertebrates [1], amniotes [2,42], turtles [43], birds [44], ray-finned fishes [45], and chipmunks [46]. Furthermore, UCEs have been shown to be an important resource for gathering information from museum specimens [47], and to be useful at shallow evolutionary time scales in birds [48].

In plant systems, genome skimming [33] has perhaps had the most impact for assembling phylogenetic datasets from HTS data. In contrast to sequence capture approaches that require preliminary genomic data for capture bait design, genome skimming requires no pre-existing genomic information. Genome skimming is a reference-guided approach that takes advantage of high copy regions in the genome, e.g., nuclear rDNA, the plastome, and the mitochondrial genome. By using reference sequences, this method ‘skims’ out targeted regions from low coverage genomic data. This approach has been used to recover the mitochondrial and chloroplast genomes in the genus *Asclepias* L. [33], study introgression in *Fragaria* L. species [49], identify horizontal transfer of DNA from the mitochondrion to the chloroplast in *Asclepias syriaca* L. [50], resolve phylogenetic relationships in the family Chrysobalanaceae [51], recover plastomes across multiple genera [23], and was also used to assemble the plastomes used for microfluidic PCR primer design in this study.

Both the UCE sequence capture and genome skimming approaches share similar technical and fundamental constraints that make their utility for phylogenetics at low-taxonomic levels with large sampling strategies limited. First, both of these methods are limited by the need to first construct HTS libraries for each sample in the study, a step that greatly increases the time and costs of the experiment. Second, variable regions flanking the UCEs are often captured at much reduced depth as one moves away from the UCE, or the UCE is lost completely if the target taxon is phylogenetically divergent from the one used in the bait design [3,48]. Smith et al. [48] found that UCEs containing variable flanking regions were usually not recovered across all samples if the variable regions extended more than 300 bp from the UCE probe. This is unfortunate, given that the more variable regions are of potentially greater utility for interspecific phylogenetic and population genetic studies. Likewise, genome skimming from low-coverage genomic data is most useful for recovering high-copy number regions in the genome; however, regions with lower representation numbers, such as single copy nuclear genes, are likely to be recovered in some samples and missed completely in others [52], depending on the depth of the low-coverage genomic data and the phylogenetic distance of the references used for mapping. Both of these cases result in the introduction of missing data, which could potentially lead to incorrect or misleading phylogenetic inferences [53]. In contrast, the large scale targeting of chloroplast, nuclear rDNA, and multiple independent single-copy nuclear genes using microfluidic PCR arrays and HTS circumvents many of these problems.

Our approach is similar in theory to targeted amplicon sequencing (TAS) methods [e.g., 37,38], but contains major improvements in efficiency. For example, Bybee et al. [37] implemented a first round of PCRs to amply each target region, which was then reamplified in a second round of PCRs to incorporate barcodes and HTS adapters. While using this two-reaction approach allows for more flexibility in the annealing temperature of target specific primers, this approach is labor intensive and thus difficult to scale to hundreds of samples and/or a large number of targets to take full advantage of the current yield of most HTS platforms. In their study, Bybee et al. [37] amplified six genes for 44 taxa from Pancrustacea, which translates to performing 12 PCRs for each of the 44 taxa to amplify and tag each amplicon. At this scale, both in terms of the number of samples and the number of loci, this method may be more favorable than the approach proposed here, however, once 48 or 96 different primer pairs are used to amplify hundreds of samples, this method becomes inefficient. We believe that experiments with high numbers of samples and loci are quickly becoming more common, and that the fields of systematics, phylogenetics, and population biology need the tools to deal with this type of sampling.

In this paper, we test the performance and utility of our targeted, subgenomic approach using the Neotropical clade of the plant genus *Bartsia* L. (Orobanchaceae) (Fig. 1). This clade is comprised of approximately 45 closely related species that are part of an ongoing rapid and recent radiation in the páramo ecosystem above tree line (∼2900m in elevation) throughout the Andes [54]. Using minimal genomic resources collected via plastome sequencing [23] and low coverage genome sequencing in representative species of *Bartsia*, we present an approach for designing microfluidic PCR primer combinations for amplifying i) the most variable regions of the plastome (referred to as the chloroplast set henceforth), ii) the commonly sequenced ITS and ETS regions of the nuclear rDNA repeat, and iii) a suite of putatively single-copy nuclear loci (ii and iii are referred to as the nuclear set henceforth). Our targeted subgenomic approach generated a large multilocus dataset across hundreds of samples, which allowed us to investigate evolutionary relationships at the species level. While shotgun approaches yield more data, the great majority of these data are highly conserved across samples and thus phylogenetically uninformative. By focusing on targeted loci and not whole genomes, we were able to maximize the yield of shared and phylogenetically informative data across a significantly greater number of samples, which is ideal for phylogenetic studies at low taxonomic levels.

## Methods

### Microfluidics PCR Primer Design and Validation

#### Preliminary data acquisition

Data used for chloroplast and nuclear microfluidics PCR primer design were generated from two taxa using long PCR to generate Plastome DNA templates for HTS [23] and three taxa (4 samples, Table 1) using genome skimming [33]. DNA was extracted from ∼0.02 g of silica gel-dried tissue using a modified 2X CTAB method [55], yielding 30 to 70 ng/μL of DNA per sample. Genomic DNAs were sheared by nebulization at 30 psi for 70 sec, yielding an average shear size of 500bp as measured by a Bioanalyzer High-Sensitivity Chip (Agilent Technologies, Inc., Santa Clara, California, USA). Sequencing libraries were constructed using the Illumina TruSeq library preparation kit and protocol (Illumina Inc., San Diego, California, USA) and were standardized at 2nM prior to sequencing. Library concentrations were determined using the KAPA qPCR kit (KK4835) (Kapa Biosystems, Woburn, Massachusetts, USA) on an ABI StepOnePlus Real-Time PCR System (Life Technologies, Grand Island, New York, USA). The long PCR libraries and one genome skimming library were sequenced on an Ilumina HiSeq 2000 at the Vincent J. Coates Genomics Sequencing Laboratory at the University of California, Berkeley, whereas the remaining genome skimming libraries were sequenced on an Illumina HiSeq 2000 at the Genomics Core Facility at the University of Oregon (Table 1). Raw reads were cleaned using SeqyClean v 1.8.10 (https://bitbucket.org/izhbannikov/seqyclean) using defaults settings.

**Table 1.** Sample and sequencing information for preliminary data acquisition.

| Species | Collector | Platform | Type of read<br>(bp) | No. clean<br>reads | Sequencing<br>Facility | Source |
| --- | --- | --- | --- | --- | --- | --- |
|  | Uribe-Convers 2010- | Illumina HiSeq |  |  |  | Uribe-Convers et al. |
| *Bartsia inaequalis* Benth. | 022 | 2000 | 100 single end | 0.93 | Berkeley | 2014 |
|  | Uribe-Convers 2010- | Illumina HiSeq |  |  |  | Uribe-Convers et al. |
| *Bartsia stricta* (Kunth) Benth. | 024 | 2000 | 100 single end | 0.9 | Berkeley | 2014 |
|  |  | Illumina HiSeq |  |  |  |  |
| *Bartsia pedicularoides* Benth. | Antonelli 574 | 2000 | 100 single end | 46.8 | Berkeley | This study |
| *Bartsia santolinifolia* (Kunth) |  | Illumina HiSeq |  |  |  |  |
| Benth. | Uribe-Convers 2010-41 | 2000 | 100 paired end | 46.4 | UO | This study |
|  |  | Illumina HiSeq |  |  |  |  |
| *Bartsia pedicularoides* Benth. | Uribe-Convers 2011-64 | 2000 | 100 paired end | 52.9 | UO | This study |
|  |  | Illumina HiSeq |  |  |  |  |
| *Bartsia serrata* Molau | Uribe-Convers 2012-15 | 2000 | 100 paired end | 65.1 | UO | This study |

Sequencing information of the samples used during the preliminary data acquisition step. Type of read refers to the length of the read in base pairs (bp), and if it was single or paired end. Number of reads denotes the number of raw reads in millions. Berkeley = Vincent J. Coates Genomics Sequencing Laboratory at the University of California, Berkeley; UO = Genomics Core Facility at the University of Oregon.

#### Chloroplast target selection

For the chloroplast PCR primer set, the cleaned reads were assembled against a reference genome (*Sesamum indicum* L., GenBank accession JN637766) using the Alignreads pipeline v. 2.25 [52], and visually inspected using Geneious R6 v6.1.5 (Biomatters, Auckland, New Zealand). Six complete plastomes—two from the long PCR approach and four from genome skimming—were aligned using MAFFT v.7.017b in its default settings [56]. To target the putatively most phylogenetically informative regions of the plastome, we developed custom scripts in R [57] to identify the most variable regions in the alignment that were flanked by conserved regions, and spanned between 400bp and 1000bp. This allowed us to rank and prioritize regions in the plastome for primer design (Scripts deposited in the Dryad Digital Repository: http://dx.doi.org/10.5061/dryad.fh592).

#### Nuclear target selection

For the nuclear PCR primer set, cleaned reads from the genome skimming samples were compared to two publicly available genomic databases, a list of the pentatricopeptide repeat genes (PPR) and the conserved orthologous set II (COSII), using the BLAST-Like Alignment Tool (BLAT, tileSize=7, minIdentity =80) [58]. We chose these two reference databases because a list of 127 PPR loci was shown to have a single ortholog in both rice (*Oryza sativa* L.) and *Arabidopsis thaliana* (L.) Heynh. [59], and was previously used to successfully infer the phylogenetic relationships of the plant family Verbenaceae and the *Verbena* L. complex [60], and in the plant clade Campanuloideae (Campanulaceae) [61]. Similarly, the COSII genes have been identified to be putatively single-copy and orthologous across the Euasterid plant clade [62], and several loci have been used for phylogenetic reconstructions of closely related species in the plant families Orobanchaceae [63] and Solanaceae [64].

Using a custom R-script (deposited in the Dryad Digital Repository: http://dx.doi.org/10.5061/dryad.fh592), reads that matched any reference gene from these two databases were kept, binned with their respective reference locus, and aligned using MAFFT in its default settings. We then used the online tool IntronFinder (http://solgenomics.net/, last accessed in January 2014) from the Sol Genomics Network [65] to predict exon/intron junction positions in the COSII genes. The PPR genes do not contain introns [59] and thus this step was not necessary for these loci. Reference loci that had reads from at least two taxa aligned to them forming conserved ‘islands’ separated by 400-800bp, including estimated introns with an assumed average length of 100bp, were selected for primer design (Fig. 2A). Additionally, an alignment from nuclear rDNA internal and external transcribed spacers sequences—ITS and ETS, respectively [54]—as well as an alignment of sequences of the *PHOT1* gene and one of the *PHOT2* gene [66] were made in MAFFT with default settings.

### Microfluidic PCR primer design and validation

Forward and reverse primers for the selected chloroplast regions and nuclear loci were designed using Primer3 [67-69] following the recommended criteria specified in the Fluidigm Access Array System protocol (Fluidigm, San Francisco, CA, USA), e.g., annealing temperature was set to 60°C (+/- 1°C) for all primers, and no more than three continuous nucleotides of the same base were allowed (Max Poly-X=3). Furthermore, regions identified as appropriate for primer design that were not present in every taxon in the alignment or that contained ambiguous bases (due to missing data and/or low coverage in our assemblies) were discarded. A complete list of the chosen primers can be found in S1 Table. Once the initial primer design was completed, a conserved sequence (CS) tail was added to the 5’ end of both the forward and reverse primers, CS1 and CS2 respectively (Fluidigm), resulting in the final target specific primers (TS) with universal tails (CS1-TS-F and CS2-TS-R, respectively). The purpose of the added tails (CS1 and CS2) is to provide an annealing site for the second pair of primers, which, starting from the 5’ end, are composed of the HTS adapters (e.g., PE1 or PE2 for Illumina sequencing), a sample specific forward and reverse barcode combination (e.g., BC1 and BC2), and the complementary CS sequence (CS1’ or CS2’; Fig. 2B and 2C). To avoid confusion, the first pair of primers with universal tails (CS1-TS-F and CS2-TS-R) will be referred to as the ‘target specific primers’, whereas the second pair of primers—with complementary universal tails, barcodes, and HTS adapters; PE1-BC-CS1’ and PE2-BC-CS2’—will be referred to as the ‘barcoded primers’. The CS1 and CS2 sequences were obtained from the Fluidigm Access Array System protocol, whereas the barcoded primers were custom designed to allow for dual barcoding, in order to dramatically increases the number of samples that can be multiplexed in one sequencing run (S2 Table).

### Primer validation

Due to the complexity of simultaneously using two sets of primers in one PCR, it is necessary to validate each set of primers prior to the actual microfluidic PCR amplification. Primer validation is a crucial step to ensure that no primer dimers are formed and that no interaction and/or competition between the barcoded and target specific primer pairs are negatively affecting the amplification. Primer validation was performed for each primer combination in 10 μL reactions in an Eppendorf Mastercycler ep thermocycler, following the Fluidigm Access Array System protocol. Validation reactions were performed on three species of *Bartsia* (*B. mutica* (Kunth) Benth., *B. crisafullii* N. H. Holmgren, and *B. melampyroides* (Kunth) Benth.), which represent the morphological and geographical diversity in the genus, and a negative control (using water instead of DNA), and included the following: 1μL of 10X FastStart High Fidelity Reaction Buffer without MgCl_2_ (Roche Diagnostic Corp., Indianapolis, Indiana, USA), 1.8 μL of 25 mM MgCl_2_ (Roche), 0.5 μL DMSO (Roche), 0.2 μL 10mM PCR Grade Nucleotide Mix (Roche), 0.1 μL of 5 U/μL FastStart High Fidelity Enzyme Blend (Roche), 0.5 μL of 20X Access Array Loading Reagent (Fluidigm), 2μL of 2 μM barcoded primers, 2μL of 50nM target specific primers, 0.5 μL of 30-70 ng/μL genomic DNA, 1.4 μL of PCR Certified Water (Teknova, Hollister, California, USA). Resulting amplicons from these reactions were analyzed in a QIAxcel Advance System (Qiagen, Valencia, California, USA), and primer pairs that produced a single amplicon and had no (or minimal) primer dimers were selected (Fig. 2D, S1 Table).

**Fig.2.**
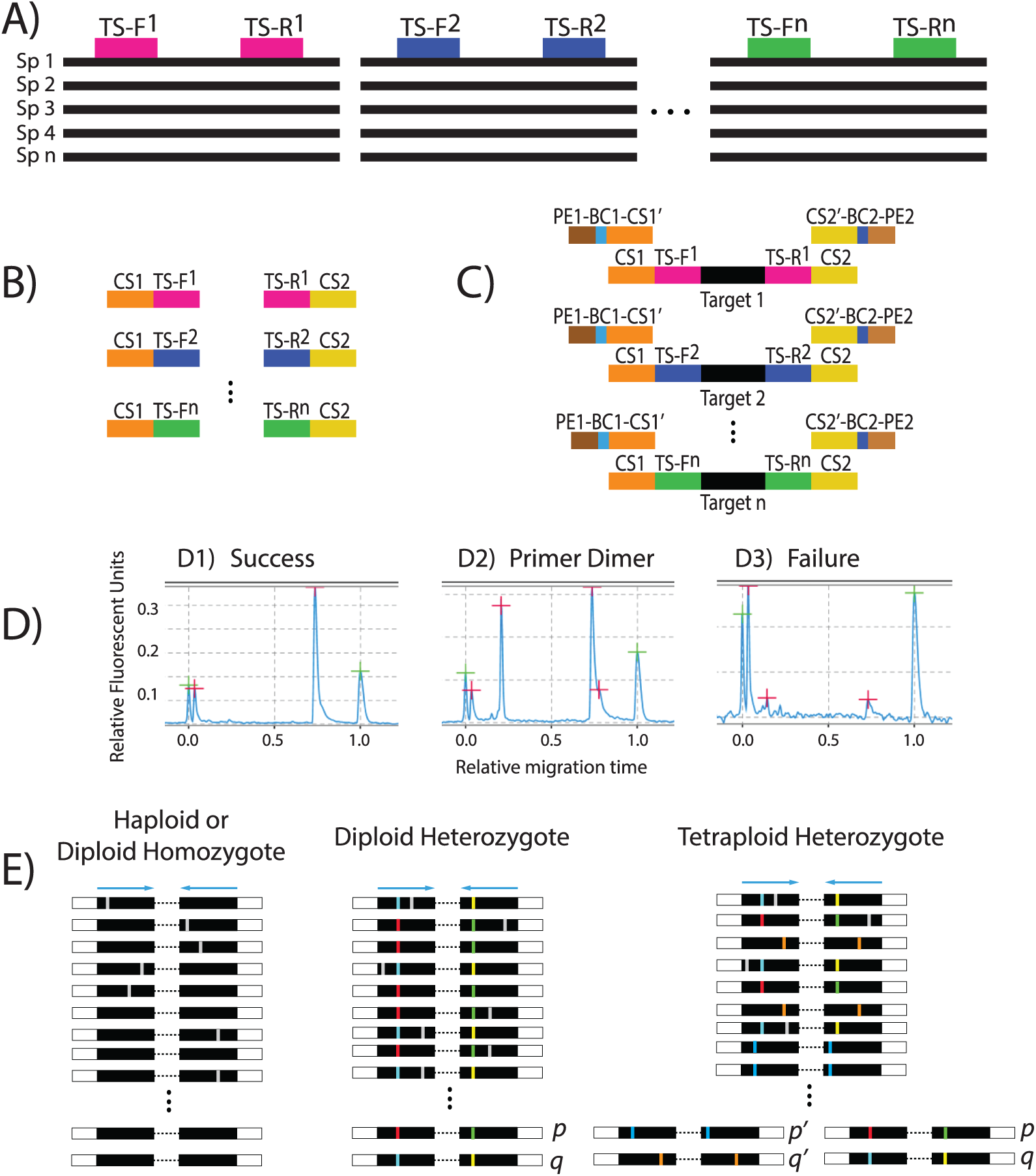
Flowchart describing the method used in this study.

Forward and reverse target specific primer combinations (TS-F and TS-R, respectively) designed in Primer3 from a multiple sequence alignment of existing genomic resources obtained in the preliminary data acquisition step. (B) The conserved sequences (CS1 and CS2) are added to the target specific primers at the time of synthesis. (C) Each target specific combination needs to be validated to ensure amplification. This is performed by simulating the microfluidic amplification reaction in a standard thermocycler, with both the first and the second pair of primers added (4-primer reaction). The second pair of primers is comprised of the sequencing adapters (e.g., PE1 and PE2, for Illumina), a barcode combination (BC) specific for each sample, and the reverse complement of the conserved sequences (CS1’ and CS2’). (D) Each reaction is analyzed for successful amplifications (D1), primer dimers (D2), or failed amplifications (D3). Only primer combinations with successful amplification and no primer dimers are chosen. (E) After sequencing, the reads are demultiplexed, sample-specific pools of amplicon sequences are generated, and groups of identical reads are identified in each pool. Pools of identical sequence reads represented by at least 5 reads and representing at least 5% of the total reads for that amplicon/sample are kept as alleles. Three examples are shown: a haploid or diploid homozygote sample with just one sequence, a diploid heterozygote sample with two different sequences (p & q), and a tetraploid heterozygote sample with two sets of homeologs (p & q and p’ & q’).

### Sampling, microfluidic PCR and sequencing

We were interested in generating data to investigate the evolutionary history of the Neotropical *Bartsia* clade [54], and thus, we sampled the complete species richness of the group, including multiple individuals per species, and some of its close relatives. A total of 74 species were represented across the 576 samples. These samples encompassed the entire geographic breadth of the South American clade, with samples ranging from northern Colombia to southern Chile (S3 Table, Fig. 3). The majority of samples were collected in the field, dried in silica-gel desiccant, and stored in airtight bags. When field-collected tissue was not available, leaf tissue was sampled from herbarium specimens (S3 Table). For all samples, DNA was extracted from ∼0.02 g of silica gel-dried tissue using a modified 2X CTAB method [55], yielding ∼30 to 70 ng/μl of DNA per sample.

**Fig.3.**
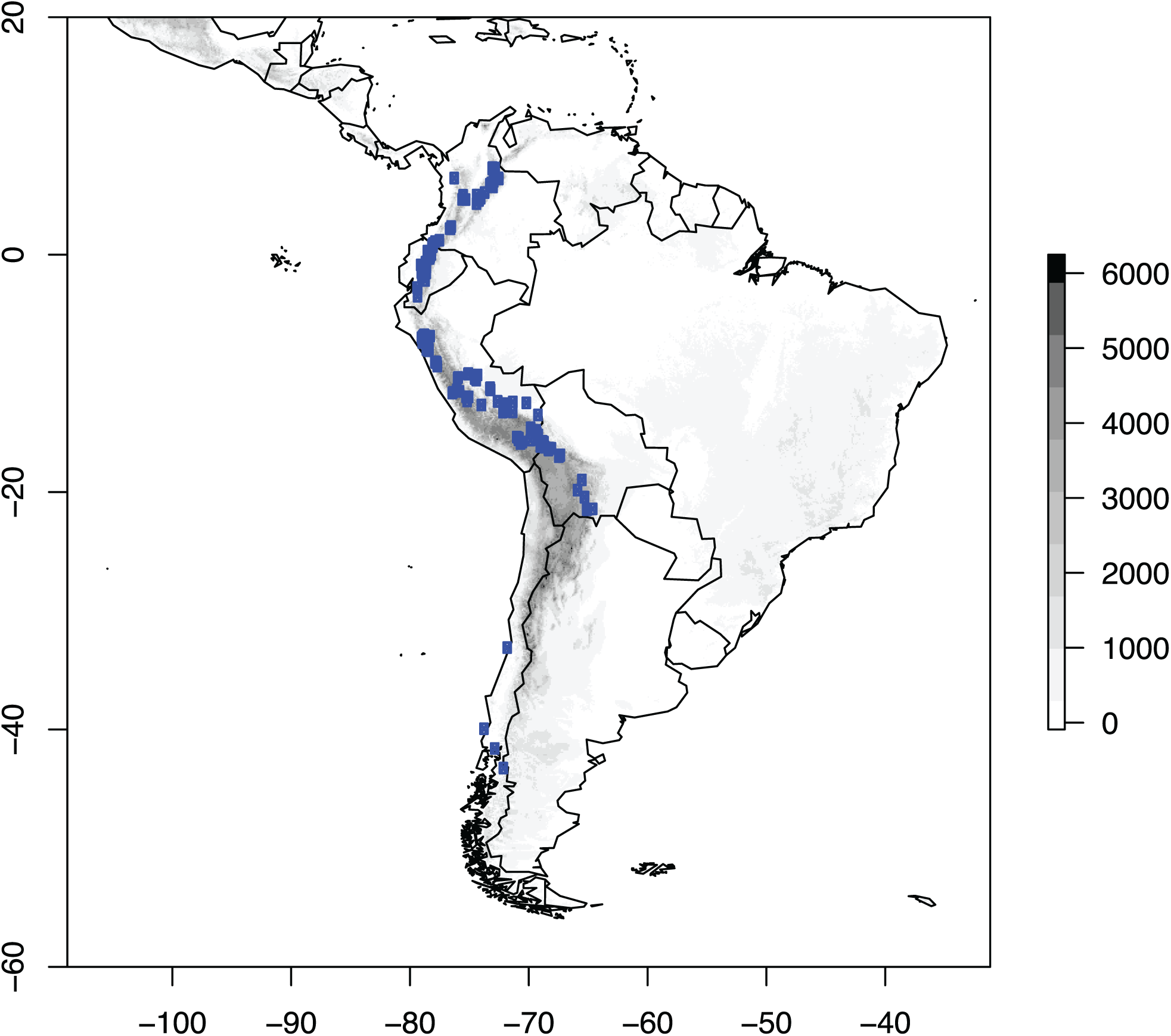
Sampling effort in the South American *Bartsia* clade.

A total of 576 samples, localities represented by dots, were collected for this study. Samples of *Bellardia trixago* L., *Bellardia viscosa* (L.) Fisch. & C.A. Mey, and *Parentucellia latifolia* (L.) Caruel not shown. The Y and X axis represent latitude and longitude, respectively, and the gray scale to the right denotes elevation in meters.

Microfluidic PCR was performed in an Access Array System (Fluidigm) using 24 (12 for the chloroplast and 12 for the nuclear set) 48.48 Access Array Integrated Fluidic Circuits (Fluidigm) following the manufacturer’s protocols. This particular array allows for 48 samples to be simultaneously amplified across 48 distinct primer pairs, resulting in 2,304 isolated and unique PCR amplicons per array. While we chose here to amplify our 48 chloroplast and 48 nuclear targets in separate microfluidic arrays, we have also had success with multiplexing genomically divergent regions such as these and performing amplification of all 96 primer pairs in a single array (i.e., target specific primers for one chloroplast region and one nuclear locus pooled prior to amplification). The amplicons were harvested from each array as per the Fluidigm Access Array System protocol and pooled per sample in equal volumes. To remove unused reagents and/or undetected primer dimers smaller than 350bp, each pool was purified with 0.6X AMPure XP beads (AgencourtT, Beverly, Massachusetts, USA). The purified pools were analyzed in a Bioanalyzer High-Sensitivity Chip (Agilent Technologies, Santa Clara, California, USA) and standardized at 13 pM using the KAPA qPCR kit (KK4835; Kapa Biosystems, Woburn, Massachusetts, USA) on an ABI StepOnePlus Real-Time PCR System (Life Technologies, Grand Island, New York, USA). The resulting pools were multiplexed in an Illumina MiSeq using the Reagent Kit version 3 with a final yield of 21.4 million 300bp paired end reads. Microfluidic PCR, downstream quality control and assurance, and Illumina sequencing was performed in the University of Idaho Institute for Bioinformatics and Evolutionary Studies Genomics Resources Core facility.

### Illumina sequence data processing

Reads from the Illumina MiSeq run were demultiplexed for sample, using the sample specific dual barcode combinations, and chloroplast region or nuclear locus, using the target specific primers with the python preprocessing application dbcAmplicons (https://github.com/msettles/dbcAmplicons). Representative sequences for each amplicon were then identified using the reduce_amplicons R script (https://github.com/msettles/dbcAmplicons/blob/master/scripts/R/reduce_amplicons.R). This pipeline allows for an optional trimming step, where the number of trimmed bases can be set independently for both the forward and reverse reads (using the “--trim-1 and --trim-2” flags in the reduce_amplicons R script). This step is especially useful in instances of high sequencing errors rates towards the end of a read, especially in read 2 on the MiSeq platform. In this study we trimmed 75 bp and 150 bp of our forward and reverse reads, respectively.

To maximize the number of amplicons recovered for each sample and each DNA region, the dbcAmplicons application allows for target specific primer matching errors less than or equal to four, as long as the final four bases of the 3’ end of the target specific primers were exact matches, thus yielding firm ends. The target specific primer sequence is removed and paired reads that overlap by at least 10 bp are joined into a single continuous sequence. Finally, for every sample at every locus, the reduce_amplicons R script produces consensus sequences (using the “-p” flag in the reduce_amplicons R script) with IUPAC ambiguity codes (-p ambiguity) for individual sites represented by more than one base when each variant is present in at least 5 reads and 5% of the total number of reads (these thresholds are adjustable using the “-s” and “–f” flags, respectively), or without ambiguities (“-p consensus”). For allele recovery (“-p occurrence”), reads are reduced to counts of identical pairs or joined paired reads. If the count represents at least 5% of the total number of reads and contains at least 5 reads (again, adjustable by the user), that amplicon is retained as a candidate allele for the sample and target (Fig. 2E). Conversely, if both the sequencing depth and minimum total percentage thresholds are not met, the sample specific target is discarded. Our allele-recovering method is based on the assumption that sequencing errors are mostly random and that reads containing errors will be represented at much lower frequencies than actual biological variation.

Because the chloroplast genome is haploid, consensus sequences were generated for each of the 48 regions (using the “-p consensus” flag in the reduce_amplicons R script). In cases where the read 1 and read 2 consensus sequences did not overlap, the reads were concatenated into a single continuous sequence. Each region was aligned with MUSCLE v3.8.31 [70] in its default settings, alignments were cleaned with Phyutility v2.2.4 [71] at a 50 percent threshold to minimize missing data due to ambiguous alignment sites, visually inspected in Geneious R6 v6.1.5 (Biomatters), and any misaligned or ambiguous sequences were discarded. Finally, the 48 chloroplast alignments were concatenated with Phyutility into a single locus.

To accommodate putative heterozygosity at nuclear loci, we generated consensus sequences with IUPAC ambiguity codes (using the “-p ambiguity” flag in the reduce_amplicons R script) for every sample at every locus, as well as individual allelic occurrences for each sample when applicable (using the “-p occurrence” flag in the reduce_amplicons R script). As with the chloroplast set, paired reads that did not overlap were concatenated, and each region was independently aligned using MUSCLE and cleaned with Phyutility at a 50 percent threshold. Because the ITS and ETS regions are physically linked in the rDNA repeat, theses regions were concatenated and treated as a single locus using Phyutility.

This data processing pipeline resulted in alignments for 47 independent nuclear loci – with allelic information, when relevant – and a concatenated chloroplast dataset. To compare alternative strategies for phylogenetic analyses, consensus sequences with IUPAC ambiguity codes for each of the 47 nuclear loci were concatenated into a single dataset (∼13,500bp; the concatenated nuclear dataset), and the concatenated nuclear dataset and concatenated chloroplast dataset were combined into a single alignment of more than 40,500bp after cleaning.

### Phylogenetic analyses

The concatenated chloroplast dataset was analyzed with PartitionFinder [72] to find the best partitioning scheme while also identifying the best-fit model of sequence evolution for each possible partition. Using these partitioning schemes and models of sequence evolution, we conducted maximum likelihood (ML) analyses as implemented in GARLI v2.0.1019 [73] with ten independent runs, each with 50 nonparametric bootstrap replicates. Bootstrap support was assessed with the program SumTrees v3.3.1 of the DendroPy v3.12.0 package [74]. Likewise, we analyzed the dataset in a Bayesian framework as implemented in MrBayes v3.2.1 [75] with the individual parameters unlinked across the data partitions. We ran two independent runs with four Markov chains each using default priors and heating values. Independent runs were started from a randomly generated tree and were sampled every 1000 generations. Convergence of the chains was determined by analyzing the plots of all parameters and the –lnL using Tracer v.1.5 [76]. Stationarity was assumed when all parameters values and the –lnL had stabilized; the likelihoods of independent runs were considered indistinguishable when the average standard deviation of split frequencies was < 0.001. A consensus trees was obtained using the sumt command in MrBayes.

The nuclear dataset was analyzed in multiple ways. First, we inferred individual gene trees for each locus using RAxML v.8.0.3 [77] to ensure that the each locus was indeed single copy. Second, we analyzed the concatenated nuclear dataset with RAxML with no topological restrictions. Third, a second analysis of the nuclear concatenated dataset with RAxML, but this time constraining the topology to make every species monophyletic (concatenation with monophyly constraints; CMC) [78]. Although not a formal coalescent-based species tree method, comparisons of the CMC approach to coalescent-based species tree approaches have found them comparable and potentially the least sensitive to taxonomic sampling [78]. Furthermore, the CMC approach is a much more computationally tractable approach than currently available coalescent-based species tree approaches on datasets of the size that we are analyzing here [but see 79 for a potentially scalable approach]. Finally, the combined dataset (chloroplast and nuclear loci) – with and without monophyly constraints - was analyzed with RAxML. Although we understand the importance of analyzing this type of dataset in a coalescent framework, the scope of the present study is not to infer the species tree or make systematic conclusions for the clade in question, which is the focus of ongoing and future work, but rather to demonstrate the efficacy of this targeted, subgenomic approach for generating large phylogenetic datasets using microfluidic PCR and HTS.

## Results

### Preliminary Data Acquisition

Low coverage genomes were sequenced for genome skimming from four samples representing three species of *Bartsia*: *B. pedicularoides* Benth. (two samples), *B. santolinifolia* (Kunth) Benth., and *B. serrata* Molau. *Bartsia pedicularoides 1* yielded ∼51.6 million 100 bp single-end reads (sequenced at the Vincent J. Coates Genomics Sequencing Laboratory at the University of California, Berkeley). The other three samples (sequenced at the Genomics Core Facility at the University of Oregon) yielded an average of ∼51.4 million 100bp paired-end reads per library (Table 1; GenBank Sequence Read Archive [SRA]: SRR2045582, SRR2045585, SRR2045588, SRR2045589). Seqyclean v 1.8.10 (https://bitbucket.org/izhbannikov/seqyclean) processing resulted in ∼46.8 million reads for *B. pedicularoides 1*, ∼52.9 million for *B. pedicularoides 2*, ∼46.4 million for *B. santolinifolia*, and ∼65.1 million for *B. serrata*. The plastomes assembled with the Alignreads pipeline from these samples had an average sequencing depth of 995x (Dryad Digital Repository: http://dx.doi.org/10.5061/dryad.fh592).

### Chloroplast and Nuclear Primer Design and Validation

The *Bartsia* chloroplast alignment of six plastomes (including only one copy of the inverted repeat) had a length of ∼125kb (Dryad Digital Repository: http://dx.doi.org/10.5061/dryad.fh592). From this alignment, we were able to design a total of 74 primer pairs that spanned the entire plastome. Following primer validation, 53 primer pairs (72% success rate) passed the validation criteria. From these, a final set of the most variable 48 primer combinations was chosen, with an average variability of 2.7% (0.8% – 7.5%) (S1 Table).

For the nuclear set, we identified 51 PPR and 762 COSII loci that matched our criteria for further primer design (i.e., enough reads matching from low-coverage genomic data to attempt primer design). The nuclear rDNA, *PHOT1*, and *PHOT2* alignments (aligned lengths of 6,711bp, 578 bp, 1,272bp - Dryad Digital Repository: http://dx.doi.org/10.5061/dryad.fh592) all contained multiple places to design primers based on our criteria. A total of 188 primer pairs were designed from all datasets (S1 Table). From those, 44 belonged to the PPR gene family, 130 to COSII, 8 to the nrDNA repeat, 3 to *PHOT1*, and 3 to *PHOT2*. After validation, 26 primer pairs were chosen for PPR (59.1% success rate), 25 for COSII (19.2 % success rate), 7 for the nuclear rDNA (87.5 % success rate), 0 for *PHOT1* (0 % success rate), and 3 for *PHOT2* (100 % success rate). Finally, the primer pair amplifying the longest target sequence was chosen among the various possibilities for the nuclear rDNA and *PHOT2* loci.

### Sampling, Microfluidic PCR and Sequencing

To fully capture the morphological, genetic, and geographical diversity of the South American *Bartsia* clade, and to demonstrate the efficiency of this approach for molecular phylogenetic studies at low-taxonomic levels, we included 576 samples (S3 Table) that represented 46 species of the clade and 28 related taxa as outgroups (included primarily to evaluate how far outside of the target group primers would successfully amplify targeted loci). Microfluidic amplification of the samples using 24 48.48 Access Array Integrated Fluidic Circuits (Fluidigm) resulted in up to 96 amplicons per sample (a total of 55,296 microfluidic reactions). After pooling and normalizing amplicons for each sample, pools were sequenced on an Illumina MiSeq platform with the Reagent Kit version 3, yielding ∼20.3 million 300 bp paired-end reads. Raw reads were deposited in the GenBank Sequence Read Archive (SRA SRP058302).

### Data Processing

Following processing with dbcAmplicons, ∼16.9 million reads (77.7%) were sufficiently matched to both barcodes (sample specific) and primers (target specific). Discarded reads (∼4.5 million reads) were a combination of PhiX Control v3 (Illumina; ∼3.2 million reads, or 15%) and reads that did not pass our criteria for matching both barcodes and the primers (∼1.3 million reads, or 7.3%).

***Chloroplast set.*** Of the 576 samples used in this study, 528 (91.7%) amplified at least one chloroplast DNA amplicon and 486 (84.0%) produced more than 40 amplicons (>21,300bp) (Fig. 4A). The majority of the samples that did not amplify efficiently belonged to outgroup taxa that are distantly related to the South American clade of *Bartsia*, suggesting that the designed primers were too specific to work efficiently outside this clade. This highlights the importance of careful primer design and validation that is in line with the taxonomic breath of the intended study. Because our primary focus was on the South American *Bartsia* clade, and primers were designed with plastomes from this clade, these results were not surprising. Following processing with reduce_amplicons, multiple sequence alignment, and cleaning, the final chloroplast dataset included 486 samples and had an aligned length of ∼25,300bp. The majority of the samples belonged to the South American *Bartsia* clade (472), with the remaining 12 samples representing the three most closely related species (*Bellardia trixago* (L.) All., *Bellardia viscosa* (L.) Fisch. & C.A. Mey, and *Parentucellia latifolia* (L.) Caruel).

**Fig.4.**
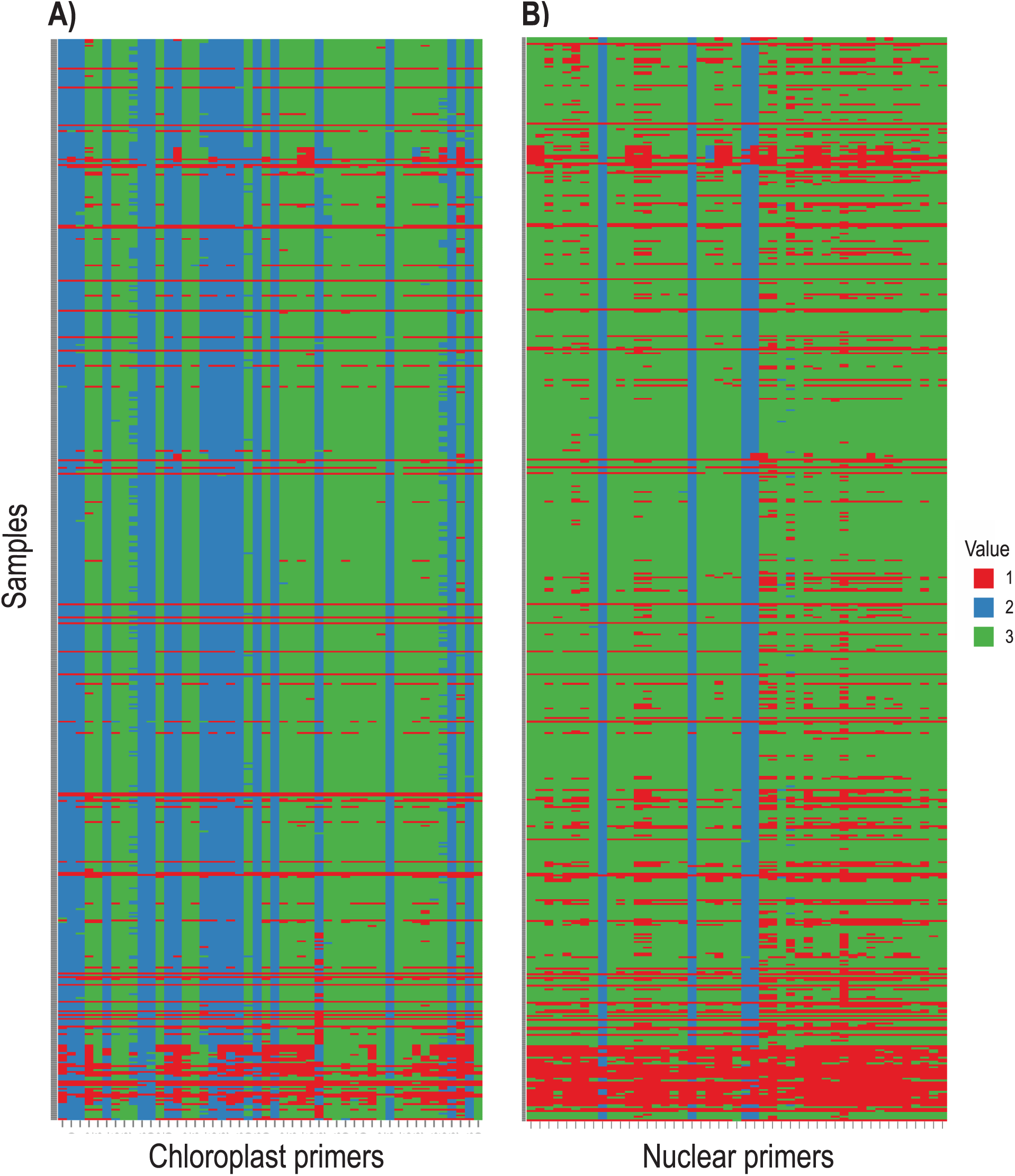
Bird’s-eye view heat map showing the target coverage for each sample

Target coverage (horizontal) for each sample (vertical) in (A) the chloroplast set and in (B) the nuclear set. These regions were processed to generate consensus sequences with ambiguities. Red indicates no amplicon was recovered either due to lack of successful amplification or mismatching of the barcodes and primers (see text for more details). Blue indicates that the forward and reverse paired reads overlapped by at least 10bp and were joined. Green indicates that the forward and reverse reads did not overlap. The group of ‘failed’ samples along the bottom of each panel are distantly-related taxa that were not included in primer design (see text).

***Nuclear set.*** For the nuclear DNA set, 47 out the 48 regions were amplified from the majority of samples (Fig. 4B), and only one region did not satisfy our amplicon demultiplexing criteria, i.e., did not meet both barcode and primer matching criteria. From these 47 regions, we were able to recover sequence data from an average of 442.7 samples (76.0%), ranging from 520 (90.3%) to 318 (55.2%). Allele recovery (from processing with reduce_amplicons) resulted in 85.41% of these samples having one allele, 13.67 % having two, 0.82% having three, and 0.10% having four alleles (S4 Table). After summarizing across every sample for each species, 17 taxa presented a combination of one and two alleles across loci (diploid as the minimum ploidy level), while the remaining 33 taxa had a combination of three and four alleles (tetraploid as the minimum ploidy level) (S4 Table). Following alignment cleaning of consensus sequences with ambiguities (from processing with reduce_amplicons), the nuclear gene regions had an average aligned length of 332bp (from 267 to 459bp). Preliminary ML phylogenetic analyses in RAxML revealed three loci where paralogous copies were amplified, and another three loci that were too variable to be unambiguously aligned (due to non-specific PCR amplification). These six loci were removed prior to downstream phylogenetic analyses, resulting in a concatenated nuclear dataset of 41 regions (nrDNA plus 40 single copy nuclear gene regions) with an aligned, cleaned length of ∼13,500bp from 363 samples. Finally, we constructed a combined concatenated matrix (nuclear and chloroplast) with an aligned, cleaned length of ∼40,500bp, including 349 samples (all sequences deposited in the Dryad Digital Repository: http://dx.doi.org/10.5061/dryad.fh592).

### Phylogenetic analyses

Individual nuclear gene trees were largely unresolved, and paralogous loci were identified in three of the 48 loci. The concatenated chloroplast dataset was first analyzed with PartitionFinder to identify the best partitioning scheme, while also selecting for the best-fit model of sequence evolution for each possible partition. This analysis resulted in 11 partitions with the following models of sequence evolution: K81uf+I+G, K81uf+I, TrN+I+G, TVM+I+G, F81, K80+I, TVMef+I+G, TVMef+I+G, F81, TVM+I+G, TVM+I+G (data partitions and corresponding models can be found in Dryad Digital Repository: http://dx.doi.org/10.5061/dryad.fh592). Analyses in ML and Bayesian frameworks, in GARLI and MrBayes, respectively, resulted in the same overall phylogenetic relationships among the samples. The same is true for the nuclear concatenated, the combined (nuclear and chloroplast), and the concatenation with monophyly constraints (CMC) analyses, which resulted in the same overall relationships among species. Because every analysis resulted in a very similar tree, the results and discussion will be based on the combined concatenated dataset, with and without constraints (all tree files deposited in the Dryad Digital Repository: http://dx.doi.org/10.5061/dryad.fh592).

Several clades that correspond to relationships between outgroup taxa and South American *Bartsia* species were recovered with 100% bootstrap support (BS) and a posterior probability (PP) of 1.0. First, all individuals included for the outgroup taxa *Bellardia trixago, Bellardia viscosa*, and *Parentucellia latifolia* were reciprocally monophyletic. Second, all accessions of South American *Bartsia* species (the large majority of the sampling in this study) were monophyletic, and *P. latifolia* is the sister group to this clade. Third, *Bellardia trixago* and *B. viscosa* formed a distinct clade, and this clade is sister to the *P. latifolia* plus South American *Bartsia* clade (Fig. 5).

**Fig.5.**
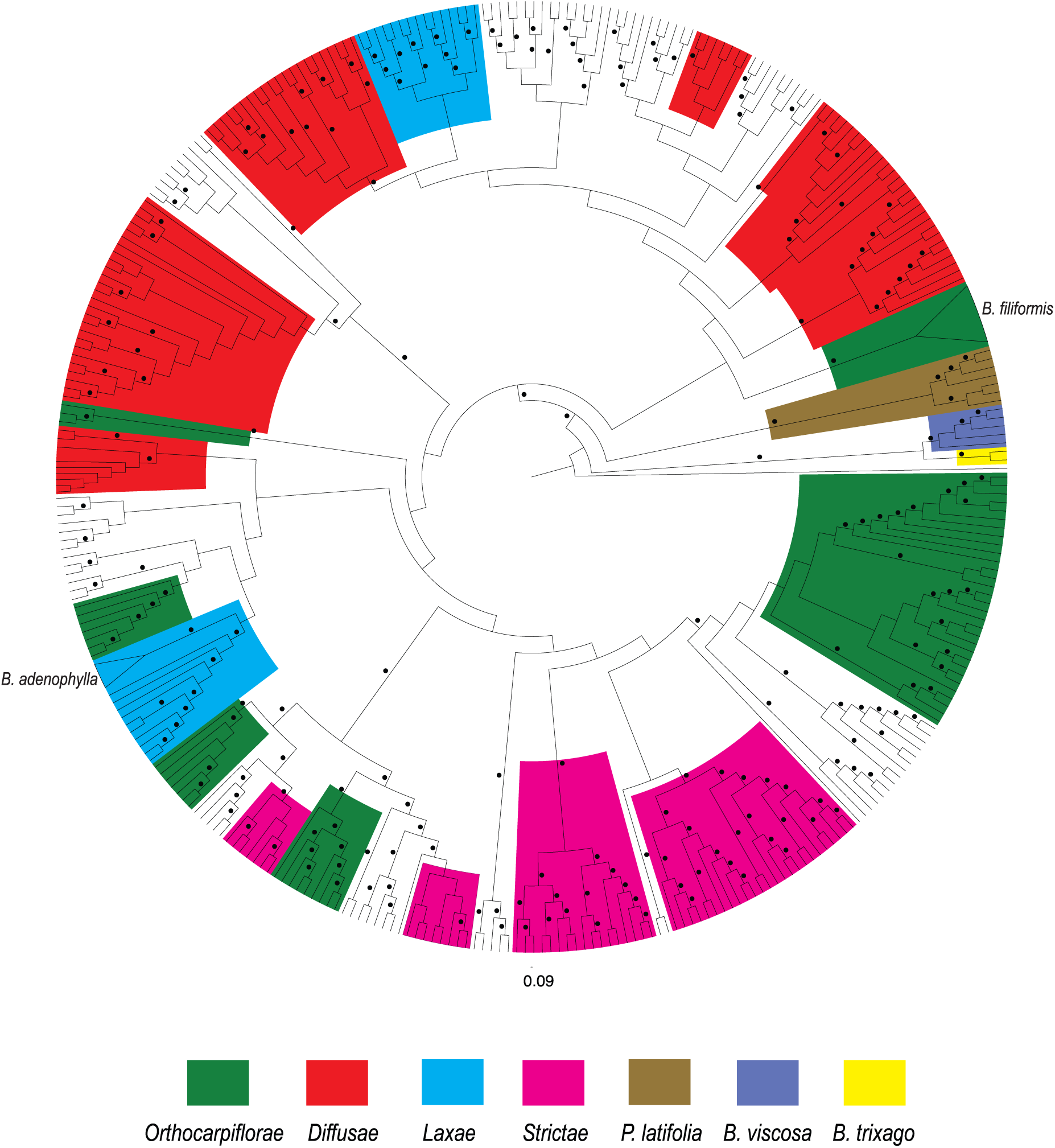
Cladogram of the phylogenetic relationships of the South American *Bartsia* species and closely related taxa

Cladogram based on the maximum likelihood analysis in RAxML on the combined (chloroplast and nuclear) unconstrained dataset. Circles above the branches represent maximum likelihood bootstrap support (BS) higher than 50. Clades containing individuals from species from only one of the four morphological sections of the South American clade [sensu 80] have been colored, as well as the three closely related taxa (*Bellardia trixago trixago* (L.) All., *Bellardia viscosa* Fisch. & C.A. Mey, and *Parentucellia latifolia* (L.) Caruel. The only three that were recovered as monophyletic are indicated on the tree and their clade collapsed in a triangle.

Support for backbone relationships in the South American *Bartsia* clade is low, and thus, very few systematic conclusions can be made at this point. First, we did not recover four monophyletic groups corresponding to the four morphological sections [sensu 80]. However, there are several clades that do contain multiple species from the same section with moderate support (Fig. 5). Furthermore, individuals of most of the species were recovered in multiple different clades, and in fact only two species (*B. filiformis* Wedd. and *B. adenophylla* Molau) were monophyletic. This is not surprising, given the fact that the South American clade has been shown to be a recent and rapid radiation [54], and processes like coalescent stochasticity, hybridization, and introgression may be playing a large role in the evolution of these taxa. It will be necessary to conduct species tree (e.g., coalescent-based) and network analyses to confidently elucidate relationships among species in this clade.

The CMC ‘pseudo-species tree’ analysis recovered most of the same clades containing species within the same taxonomic sections. Interestingly, enforcing monophyly of individuals belonging to named species reduced BS support of the backbone relationships even further (Fig. 6), indicating that sequences from some of the individuals that were constrained to be monophyletic clearly violate this assumption.

**Fig.6.**
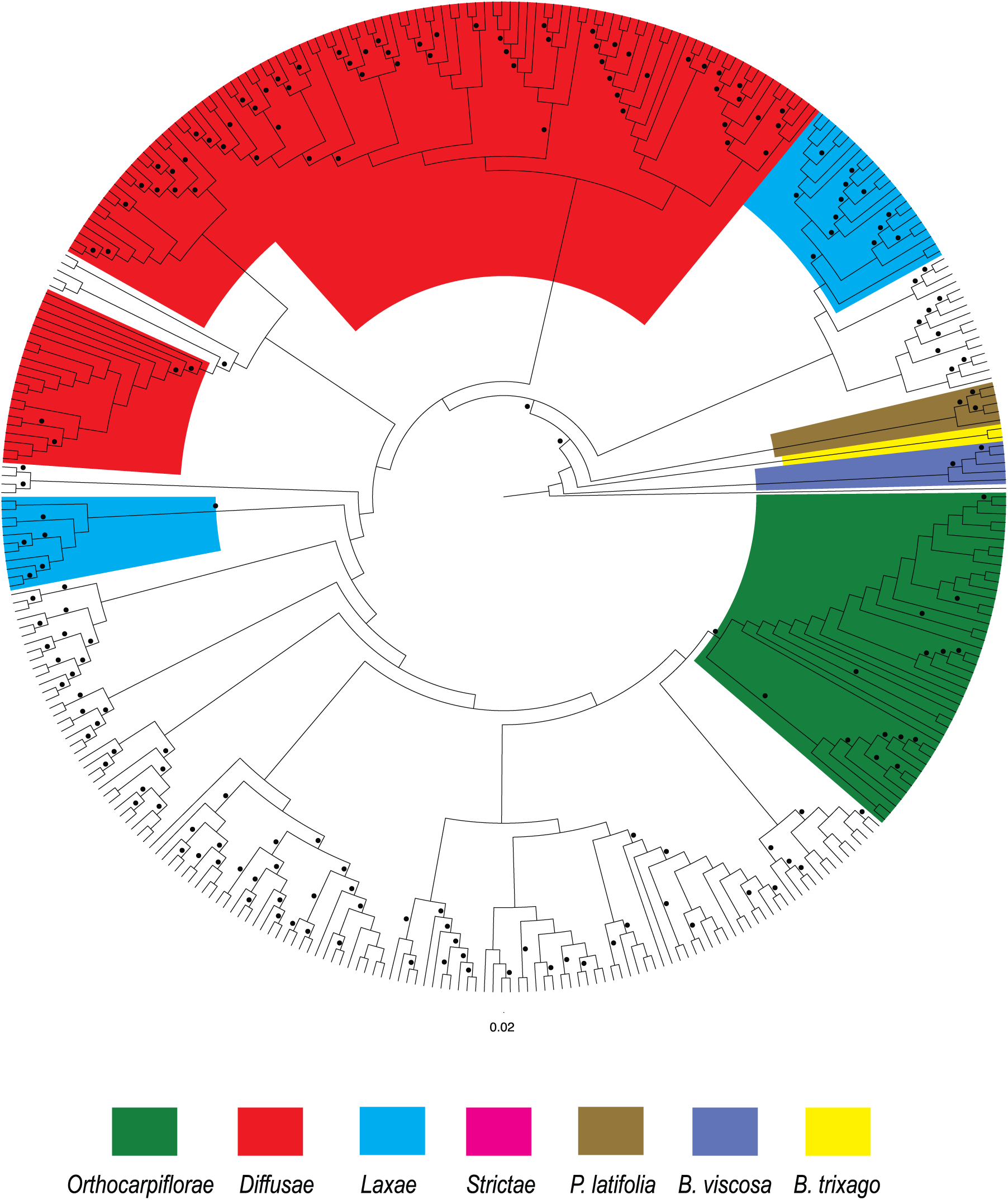
Cladogram of the phylogenetic relationships of the South American *Bartsia* species and closely related taxa

Cladogram based on the maximum likelihood analysis in RAxML on the combined (chloroplast and nuclear) dataset using the concatenation with monophyly constraints (CMC) approach on the combined dataset. Circles above the branches represent maximum likelihood bootstrap support (BS) higher than 50. Clades containing at least two species from one of the four morphological section of the South American clade [sensu 80] have been colored, as well as the three closely related taxa (*Bellardia trixago trixago* (L.) All., *Bellardia viscosa* Fisch. & C.A. Mey, and *Parentucellia latifolia* (L.) Caruel.

## Discussion

Regardless of the HTS method used, it is clear that the field of phylogenetics, and the study of evolution in general, are quickly migrating towards larger and larger molecular datasets. The ability to produce more and longer reads, as well as reduced sequencing costs and increased computing power are making this transition easier and faster. Here, we presented a novel approach to generate large multilocus, homogeneously distributed, and targeted subgenomic datasets using microfluidic PCR and HTS.

One of the main advantages of this approach is circumventing the necessity to construct expensive genomic shotgun libraries for each sample in the experiment. This step greatly increases the time and cost of any HTS approach, effectively reducing the sample size possible for any experiment. The approach presented here takes advantage of a four-primer PCR amplification to efficiently tag multiple genomic targets with sample specific barcodes and HTS adapters. By doing so, the resulting amplicons are ready to be sequenced following standard pooling and quality control. Furthermore, the use of a sample-specific dual barcoding strategy allows for a high level of multiplexing with far fewer PCR primers. Commonly used commercial barcoding kits currently offer either 96 (NEXTflex DNA Barcode kit; Bioo Scientific, Austin, Texas, USA) or 386 barcodes (Fluidigm), but we are able to multiplex up to 1,152 samples with only 72 barcoded primers (48 forward and 24 reverse). This expands the possibilities during experimental design and takes full advantage of the yield of current HTS platforms, while maintaining low upfront costs.

An additional technical advantage of our microfluidic approach is the high throughput achieved with smaller amounts of DNA, reagents, and labor. A commercially available platform – the Fluidigm Access Array System – facilitates simultaneous amplification of 48 samples with 48 distinct primer pairs (2,304 reactions) using only 15 U of *Taq* polymerase and 1μL of 30-60ng/μL genomic DNA per sample. By conducting a simultaneous four-primer reaction, one avoids the necessity of performing multiple rounds of manual PCR to incorporate barcodes and adapters – a limitation of the Targeted Amplicon Sequencing (TAS) strategy of Bybee et al. [37]. For example, following the TAS approach, to produce tagged amplicons for the 576 samples and 96 gene regions targeted in this study, it would have required >1,100 96-well plates (or >275 384-well plates) of PCR to produce the same number of barcoded amplicons. While the TAS approach does allow for more flexibility in terms of primer design, i.e., primer annealing temperatures do not need to all be the same, and it may be possible to incorporate ambiguities into primer design, to take advantage of performing PCR in plates, significant PCR optimization would also need to be performed. Nevertheless, with high levels of taxon-by-gene region samplings, TAS becomes unpractical. Using the microfluidic PCR approach to amplicon generation and tagging, only 24 microfluidic chips were necessary to amplify and tag the 55,296 amplicons. While studies with smaller sampling strategies [e.g., 37] would likely benefit from the two-reaction TAS approach, with the ever increasing sequencing read length and throughput of HTS platforms, the microfluidic PCR approach presented here allows researchers performing large phylogenetic or population genetic studies to maximize data collection using HTS techniques.

Perhaps the most significant advantage of generating phylogenetic datasets using a targeted amplicon approach is that using our ‘read frequency’ approach (Fig. 2E), it is possible to distinguish individual alleles at heterozygous nuclear loci without requiring additional assembly or mapping steps. Because targeted loci were specifically amplified one locus per reaction per sample, when paired-end sequences from each specific barcode and primer combination are identified, heterozygous loci in diploid species appear as high frequency amplicons, allowing us to straightforwardly determine both alleles. This contrasts with methods in which genomic DNA is sheared and selected for a specific length, e.g., sequence capture and GBS/RADseq, where an assembly and/or reference-based mapping strategy is necessary to compile consensus sequences. These additional steps introduce known problems associated with the large number of de novo assemblers and mappers, e.g., varying numbers of contigs and N50 using different algorithms, performance of the algorithm based on the error model of the sequencing platform used, and computational power and time [see 81 for further details]. More importantly, most phylogenomic studies that have included nuclear data generated using HTS techniques like these, have ignored the challenge that heterozygosity presents by using ambiguity coding (e.g., [1] or by selecting only one allele and discarding others e.g., [2,48]). For phylogenetic and population genetic studies using modern coalescent-based approaches, allelic information is important when reconstructing the evolutionary histories of the genes sampled, and in cases where there is a large amount of coalescent stochasticity and/or gene flow, discarding or masking allelic information may be misleading. In a population genetic study of the North American tiger salamander (*Ambystoma tigrinum* Green) species complex, O’Neill et al. [38] used a haplotype phasing strategy to computationally determine individual alleles. For statistical phasing approaches, the number of individuals per population present in a sample is a critical factor in determining how well haplotype phase can be estimated [82], and therefore may only be appropriate for the deep population-level sampling in population genetic studies such as this, but will likely not be useful for most phylogenetic studies.

Furthermore, polyploidy is common in many plant groups, as well as in select groups of insects [83], fish [84], amphibians [85], and reptiles [86], and therefore, this is an important consideration that complicates the issue of heterozygosity even more. For example, a tetraploid species may be heterozygous at both homeologous loci, and in this scenario, one would expect to identify four sets of reads with high frequencies dominating the amplicon pool. Likewise, a tetraploid may be homozygous at one homeolog and heterozygous at the other; in this case, we would expect to identify three-sets of high frequency reads dominating the amplicon pool.

Finally, for many species of plants, ploidy levels are often unknown, or variable within a species [e.g., 87], and material appropriate for determining ploidy via chromosome counts and/or flow cytometry is not available. While at any one nuclear locus, a polyploid species may or may not be heterozygous at one or more of the homeologs, by having multiple nuclear loci in one experiment, it may be possible to calculate the frequencies of alleles across all loci and not only recover individual alleles, but potentially estimate ploidy level – or at least a minimum ploidy level depending on levels of heterozygosity. In plants, this could be especially useful for evaluating hypothesized allopolyploid events, as well as the evolutionary and ecological consequences of polyploidy when these data are analyzed in a comparative phylogenetic context.

Within the South American *Bartsia* clade, only 23 of 45 species have published chromosome numbers, and seven of these have been characterized as tetraploids based on these counts (S4 Table) [80]. Likewise, chromosome counts have been published for the European *Bartsia alpina* (diploid), and the Mediterranean species *Bellardia viscosa* (tetraploid) and *Parentucellia latifolia* (tetraploid) [80]. The reduce_amplicons pipeline employed here recovered at least three alleles in one locus for six of these species, suggesting that their minimum ploidy level might be tetraploid. Although we only recovered one or two alleles in the remaining four species (suggesting that the taxa may be diploid), it is important to keep in mind that these species comprise a very recently diverged clade, and most species occur in small, isolated populations [54], where low sequence divergence, autopolyploidy, and homozygosity may mask true ploidy levels. Something to keep in mind, however, is the number of samples (individuals) per species that were recovered as having more than two alleles. For some of these species, e.g., *Bartsia camporum* Diels, *Bartsia serrata* Molau, and *Bellardia viscosa*, the majority of individuals (more than 65%) presented at least three alleles—supporting that these taxa are indeed tetraploid. On the other hand, species such as *Bartsia pyricarpa* Molau, *Bartsia pedicularoides* Benth., and *Bartsia patens* Benth.—all of which are suggested to be tetraploid based on chromosome counts (S4 Table) [80]—were recovered with most of their individuals presenting one or two alleles and only 2% to 15% of the individuals having at least three alleles. This highlights the necessity of including more than one individual from more than one population when assessing levels of ploidy. To fully investigate the utility of this approach for bioinformatically estimating ploidy levels, chromosome counts and/or flow cytometry data from these same samples would be necessary, but this was beyond the scope of this study.

Nevertheless, these results are encouraging as they open the door for future comparisons between ‘bioinformatically karyotyped’ samples and traditional ploidy estimation experiments.

A notable limitation of the microfluidic approach that we present here is the necessity to design a relatively large number of target-specific primers to fill a microfluidic array. To do this in an efficient manner, it is necessary to first have at least some genomic resources available for your clade of interest. In our case, we had both whole plastome sequences, as well as low coverage genomic data for a small, but representative, set of species. With these preliminary data in mind, we developed an effective approach for primer design that allowed us to target 1) the most variable regions of the plastome in the South American *Bartsia* clade, 2) the ITS [88] and ETS [89] regions of the nrDNA repeat that have been used extensively at the interspecific level in plants, 3) multiple, independent nuclear genes from the intronless PPR gene set developed by Yuan et al. [59] and shown to be phylogenetically informative at the family level in Verbenaceae [90] and at the subfamily level in Campanuloideae [61], and 4) intron-spanning regions from within the COSII gene set developed by Wu et al. [62] and used within Orobanchaceae [63], and the Phototropin 2 gene used at the interspecific level in *Glandularia, Junellia* and *Verbena* in the Verbenaceae [66]. By specifically targeting the variable regions of the plastome, commonly sequenced regions of the nrDNA repeat (e.g., ITS, ETS), and multiple independent nuclear loci that range from more conserved (e.g., intronless PPR genes) to rapidly evolving nuclear introns (e.g., COSII), we were able to assemble a large, multilocus, homogeneously distributed dataset with high levels of intraspecific sampling for a complete clade of recently diverged Andean plants. Although we took, and advocate, the genome skimming approach [33,52] to develop the necessary genomic resources used here for primer development, there are a growing number of publically available databases that could also be used – e.g., for plants, Phytozome (http://www.phytozome.net), One Thousand Plants Project (1KP; http://onekp.com), IntrEST [91], and Genome 10k for animals [92] – as well as a recently published bioinformatics pipeline for identifying single-copy nuclear genes and designing target specific primers for phylogenomic analyses using existing transcriptome data [93].

### The South American Bartsia Clade

This is the first time that the interspecific relationships of the species in the South American *Bartsia* clade have been studied with such deep taxonomic sampling and with so much molecular data. From our results, it is clear that in order to fully understand the evolutionary history of the clade, phylogenetic species tree methods that explicitly incorporate mechanisms that lead to gene tree-species tree discordance (e.g., coalescent stochasticity, divergence with gene flow) are needed. However, these detailed analyses are beyond the scope of this study (which is focused on data collection approaches), and therefore, the phylogenetic results presented here are preliminary. Nevertheless, the monophyly of the group is highly supported by all analyses, and is in agreement with recent biogeographic study of the clade [54]. Interspecific relationships, however, have very little support – a pattern commonly seen in rapid radiations like this – and only two species were found to be monophyletic. These two species are taxa with small and restricted geographic distributions with likely small effective population sizes. Given that the time to coalescence is directly linked to effective population size [30], it is not unexpected that individuals from these species were monophyletic in our unconstrained analyses. Conversely, when we look at species with a large geographic distributions, and larger effective population sizes (e.g., *B. pedicularoides* Benth.), we see that the individuals are recovered in multiple different groups across the tree (Fig. 5).

Enforcing the monophyly of species has been used as a ‘pseudo-species tree’ method with good results [78], and some relationships recovered here make evolutionary sense – *Bartsia sericea* Molau and *B. crisafullii* N. Holmgren were recovered as sister species (Fig. 5) with high support. Both species are extremely similar morphologically, only differing in their life history and ploidy level (perennial vs. annual, and diploid vs. tetraploid, respectively). However, in some instances, enforcing monophyly of the species reduced the BS support of deeper branches. There are several possibilities for this result, including violations of our *a priori* species designations (i.e., incorrect species delimitations, cryptic species, etc.), severe coalescent stochasticity, ancient and/or contemporary introgression, and/or hybrid speciation. Given the recent and rapid nature of this Andean diversification, each of these (or any combination of) are potential mechanisms that increase the phylogenetic complexity of this group, and incorporating these processes into species tree estimation in this clade is the focus of ongoing systematic studies using these data.

## Conclusion

We presented a novel approach to generate large multilocus phylogenomic datasets for a large number of samples and species using microfluidic PCR and HTS. This approach allows for more control in targeting informative regions of the genome to be sequenced, resulting in datasets that are tailored to address the specific questions being asked, and that are orthologous across samples. Additionally, this method is both cost effective and time efficient, as it does not require genomic shotgun libraries to be constructed for every sample, and takes full advantage of the large multiplexing capabilities of HTS platforms. As a case study, we focused on 576 samples of the South American *Bartsia* clade, amplifying and sequencing the 48 most variable regions of the chloroplast genome, as well as 48 nuclear gene regions representing a range of both coding and non-coding data. This targeted, subgenomic strategy for the collection of multilocus data for phylogenetic studies provided us with a large, but modest, set of loci that will be appropriate for sophisticated species tree inference methods (e.g., coalescent-based, networks, concordance analyses), and provided us with the first species level phylogeny for the South American *Bartsia* clade. Furthermore, the bioinformatic approaches employed here allowed for the recovery of individual alleles in heterozygote individuals (without the need for statistical phasing), and opened the door for the exploration of bioinformatic approaches to estimating ploidy levels—an important and often overlooked consideration at low taxonomic levels.

## Acknowledgements

We would like to thank Dan New, Tamara Max, and the Institute for Bioinformatics and Evolutionary Studies (IBEST - NIH/NCRR P20RR16448 and P20RR016454) at the University of Idaho for technical assistance and bioinformatic resources. Jonathan Eastman was instrumental in R scripting and data processing. Jack Sullivan, Luke J. Harmon, Eric H. Roalson, Diego F. Morales-Briones, Matthew W. Pennell, Tracy C. Peterson, and XX anonymous reviewers offered helpful comments on the manuscript.

---

**S1 Table.**
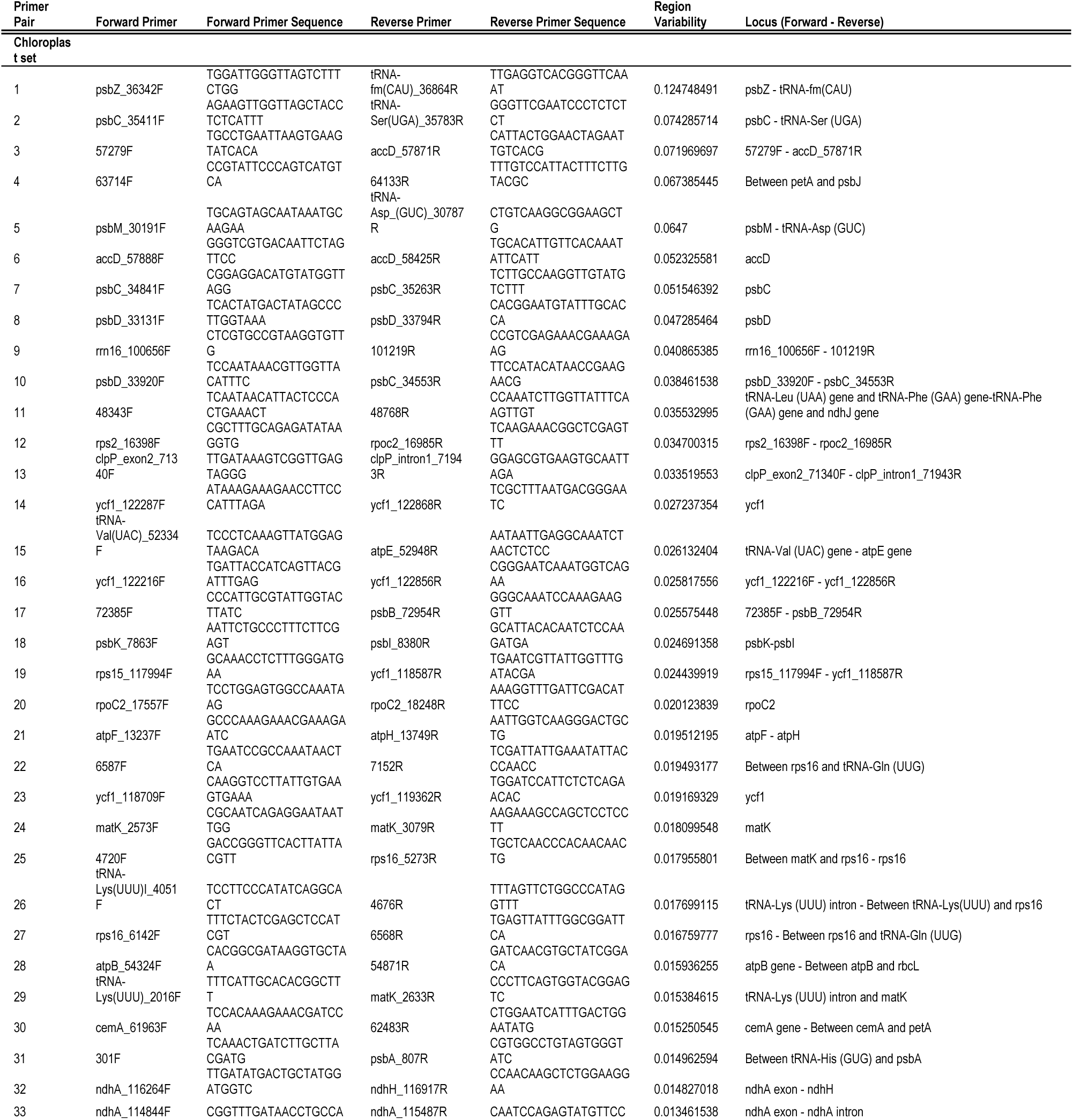
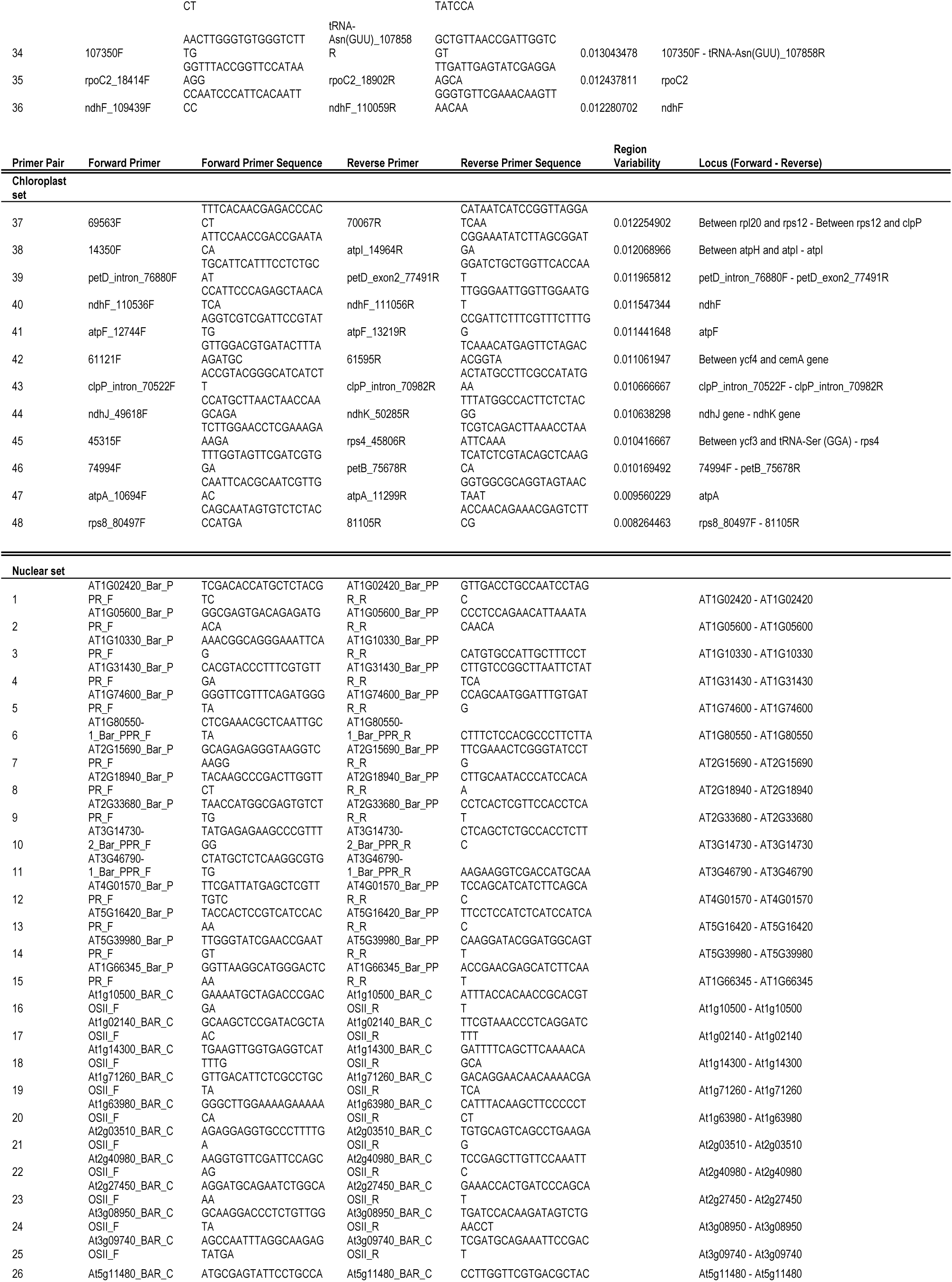
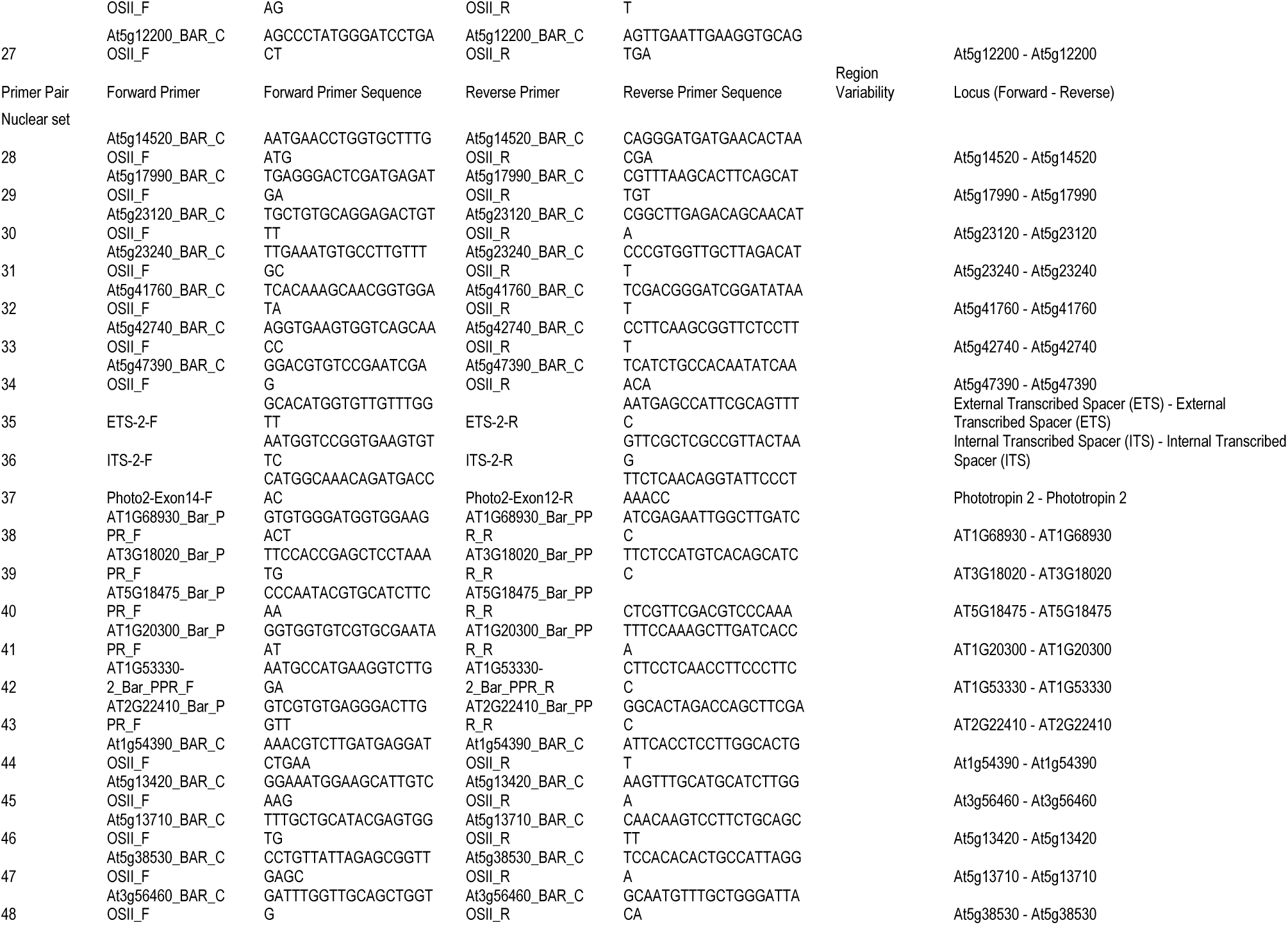
List of the primers designed and used in this study. Sequences are written in 5’-3’ direction. Variability of the region is based on substitution per site.

**S2 Table.**
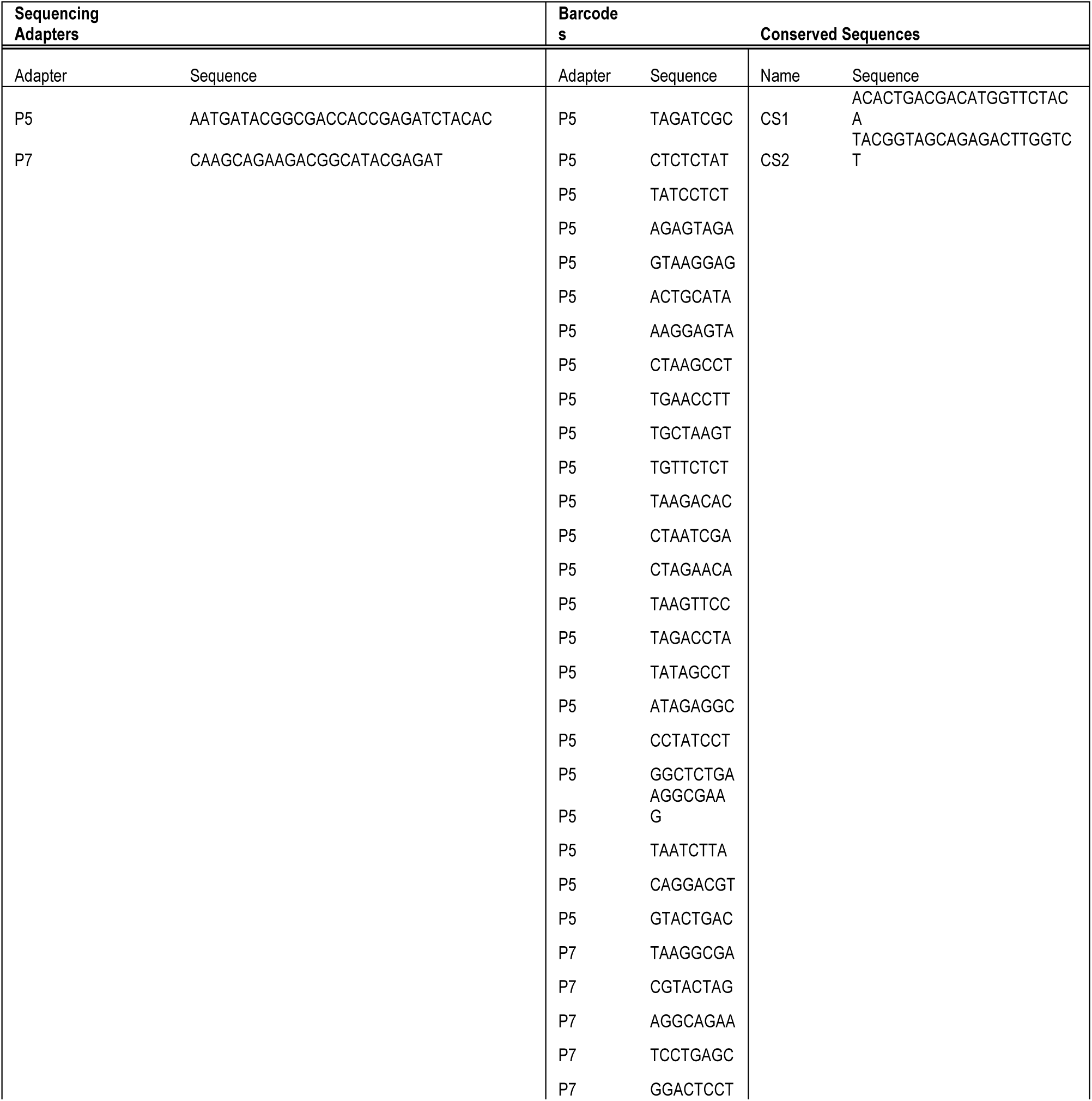
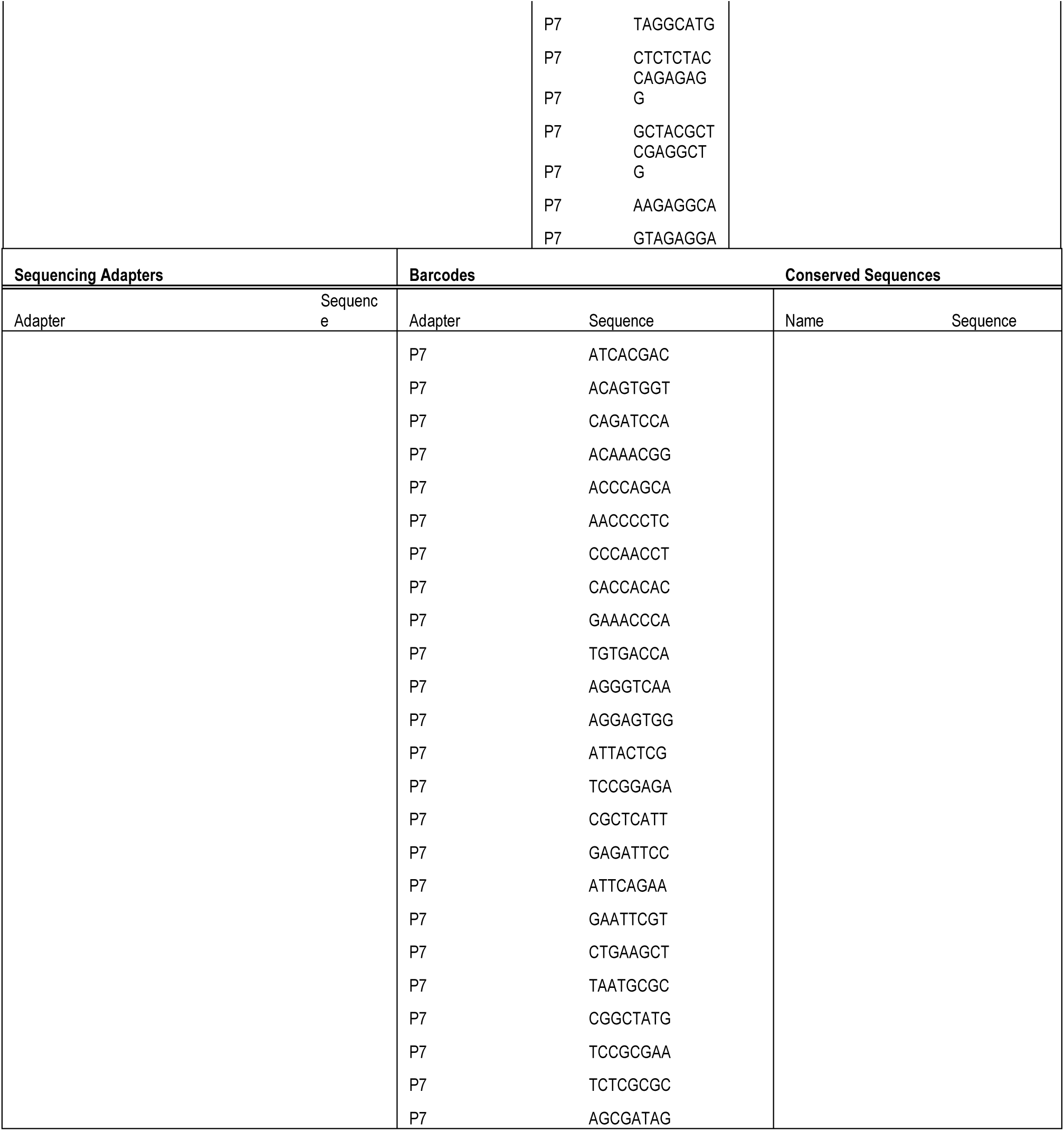
Sequences for conserved sequences 1 and 2, barcodes, and sequencing adapters. The CS1 and CS2 sequences were obtained from the Fluidigm Access Array System protocol, whereas the barcoded primers were custom designed to allow for dual barcoding, in order to dramatically increases the number of samples that can be multiplexed in one sequencing run. The sequencing adapters are the Illumina standard adapters.

**S3 Table.**
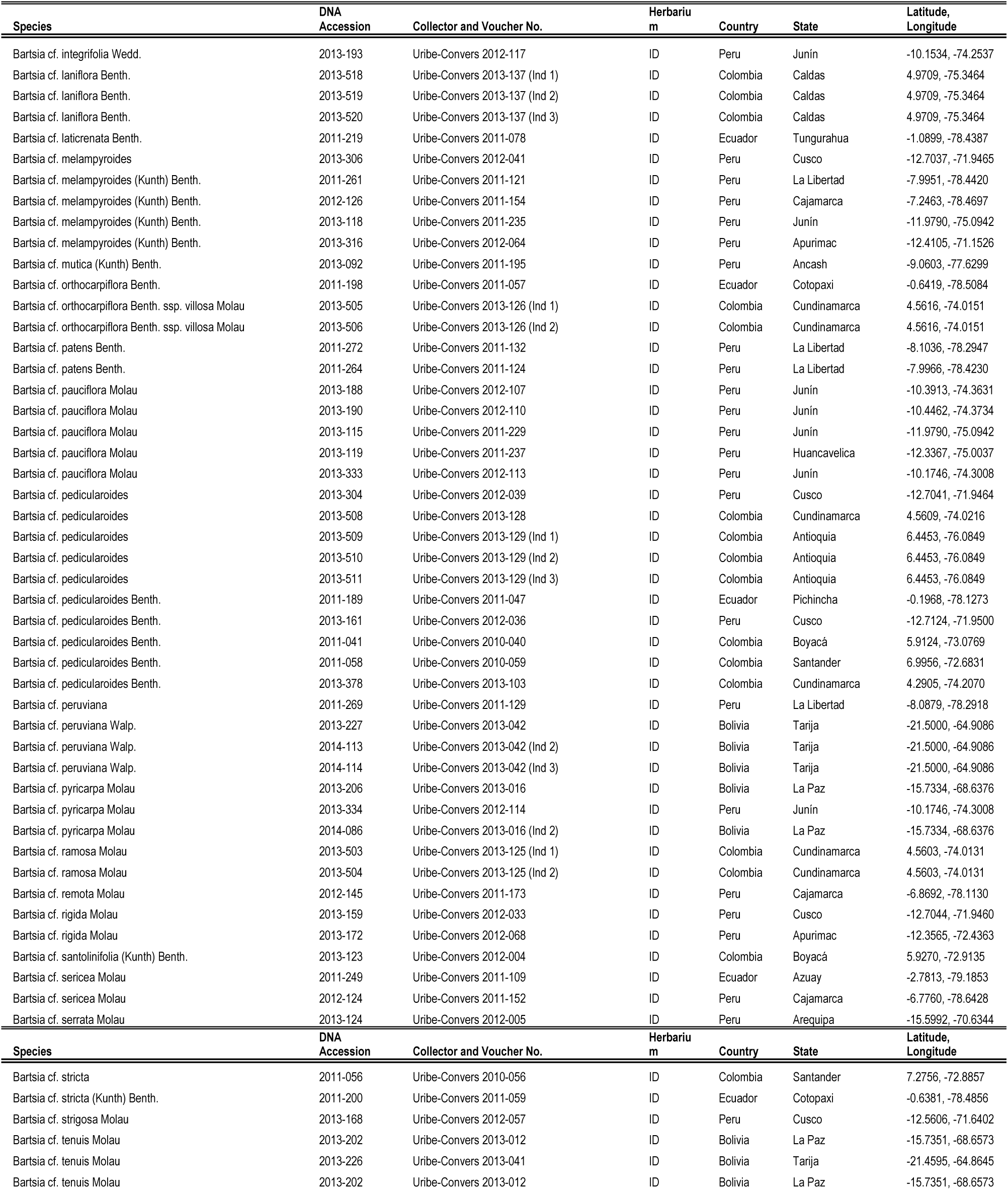
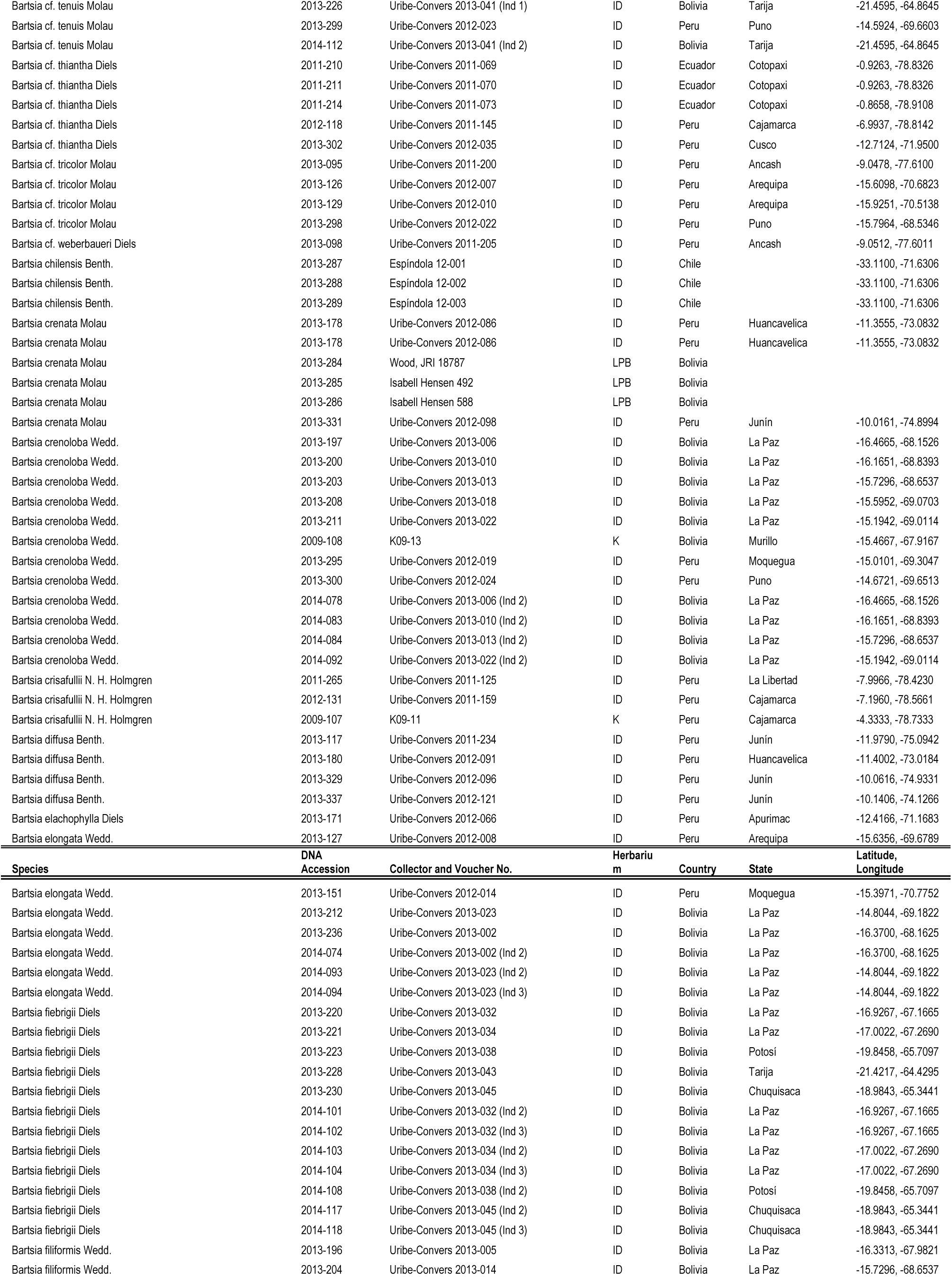
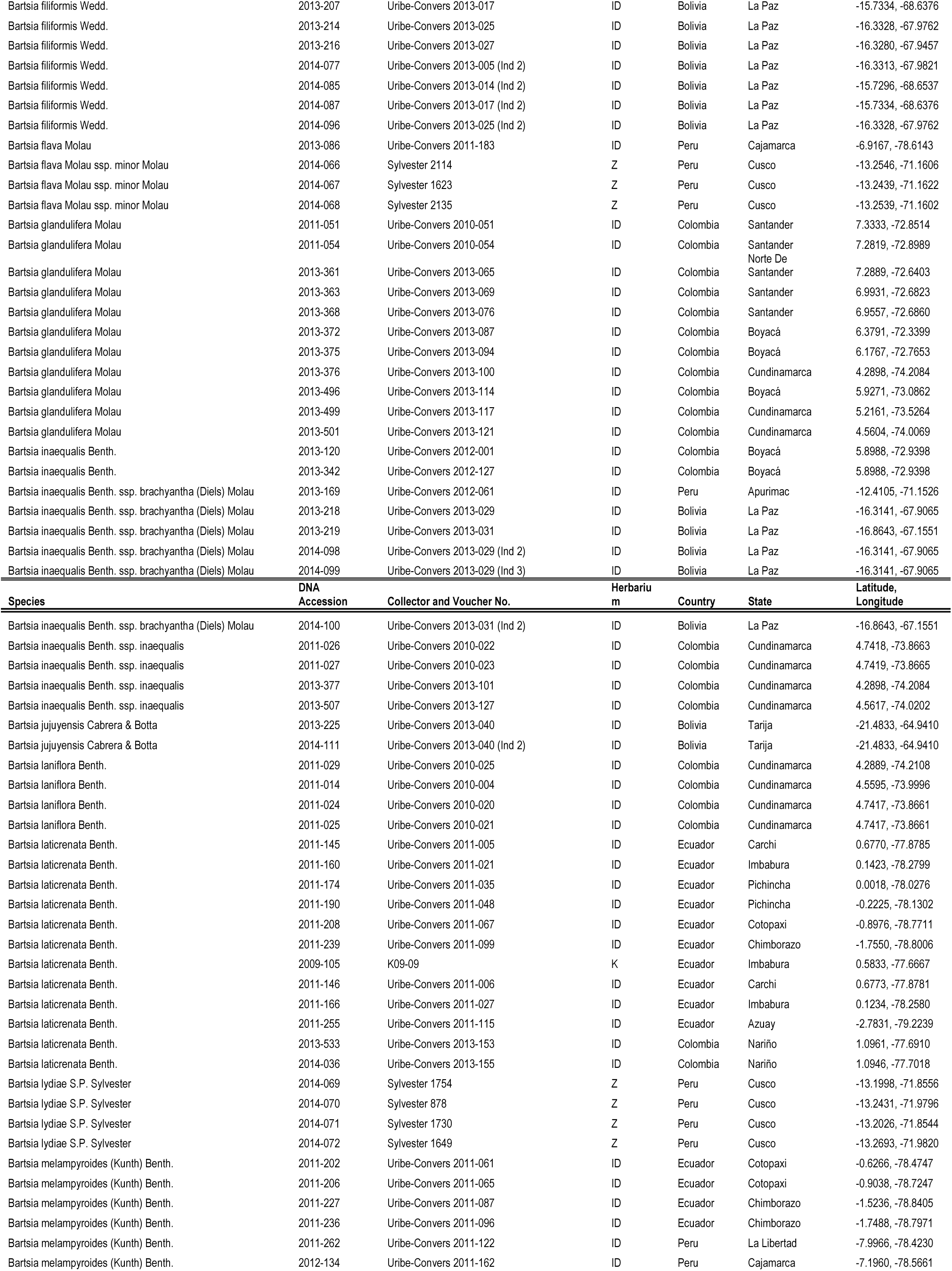
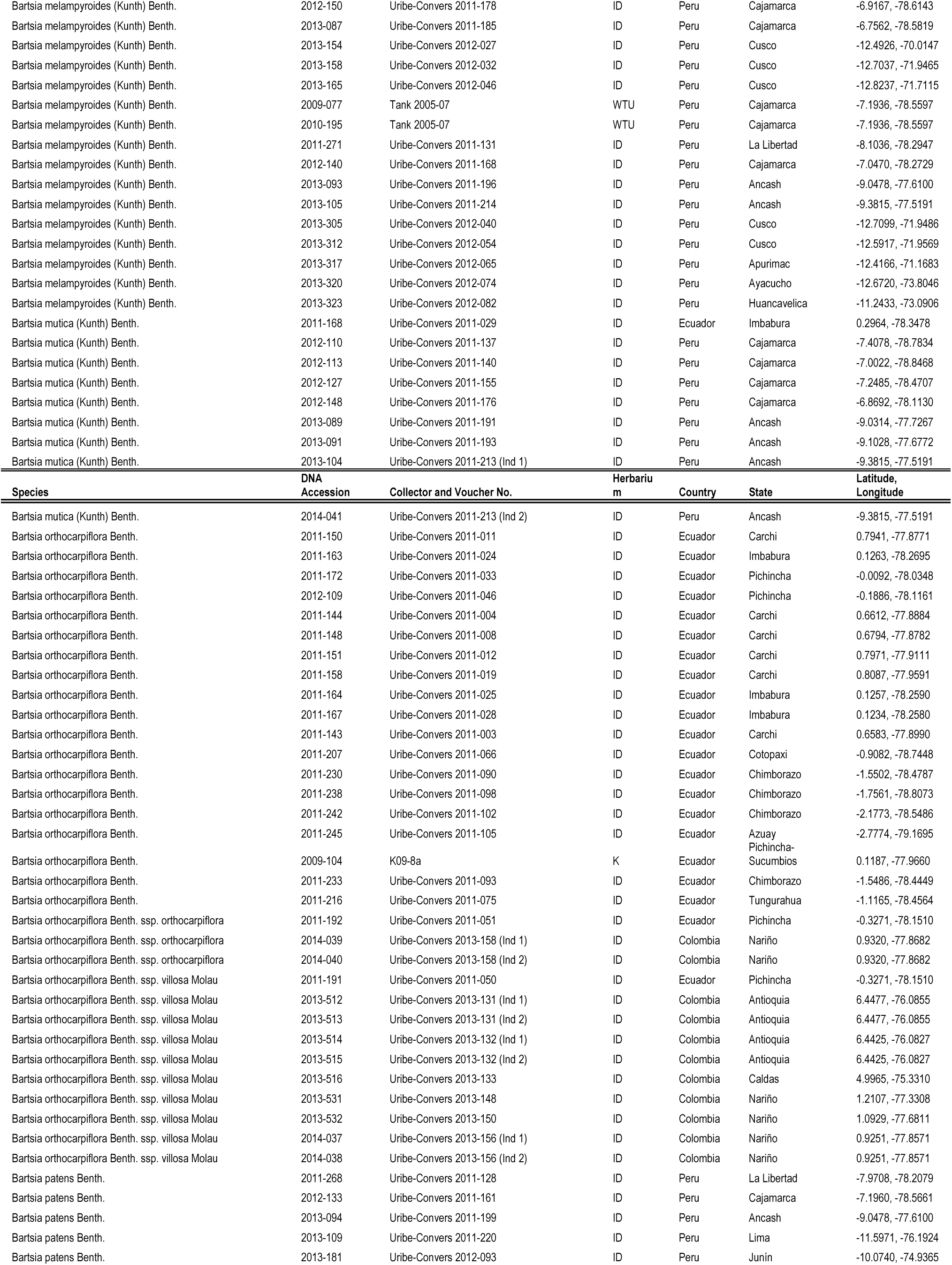
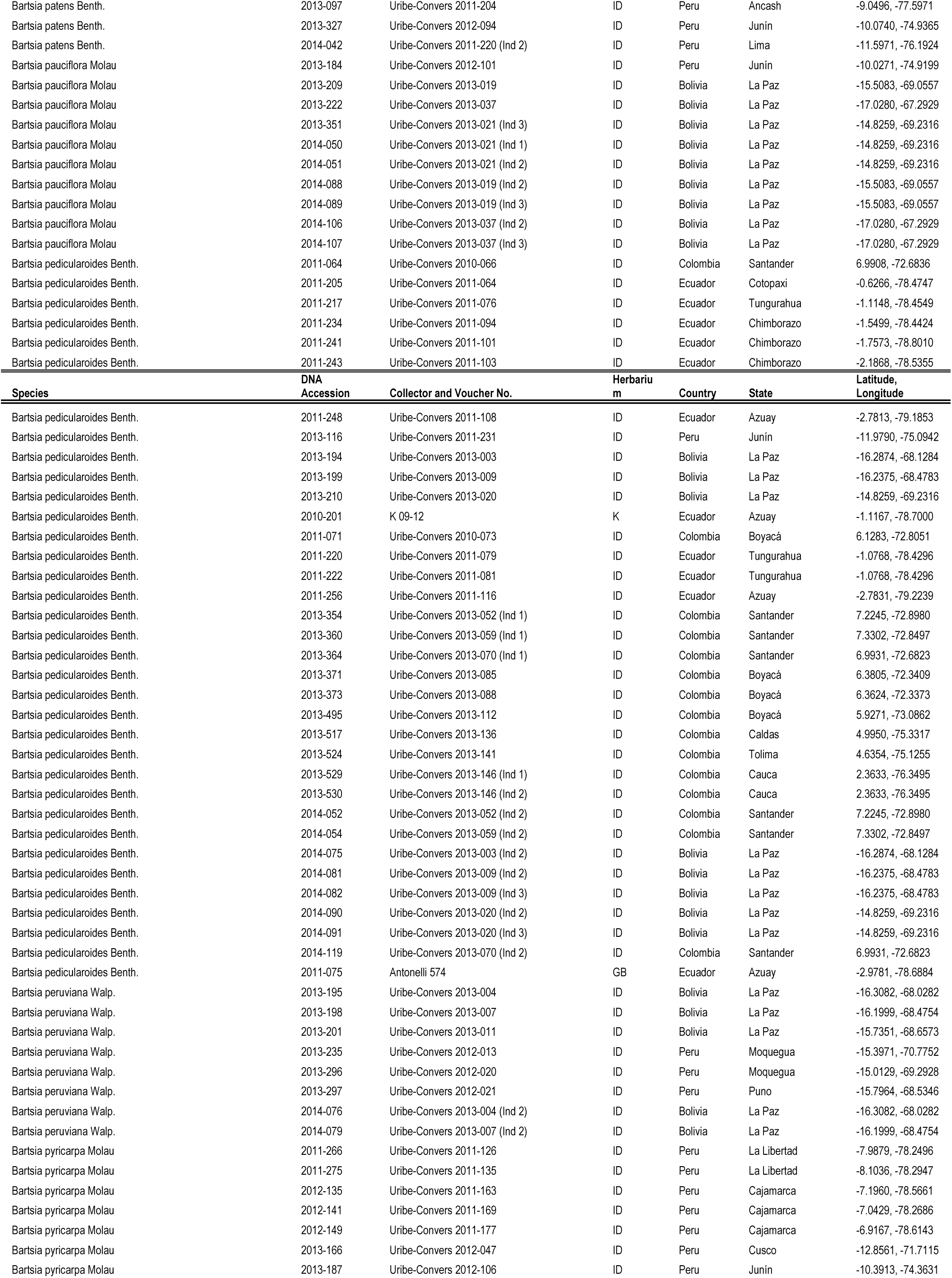
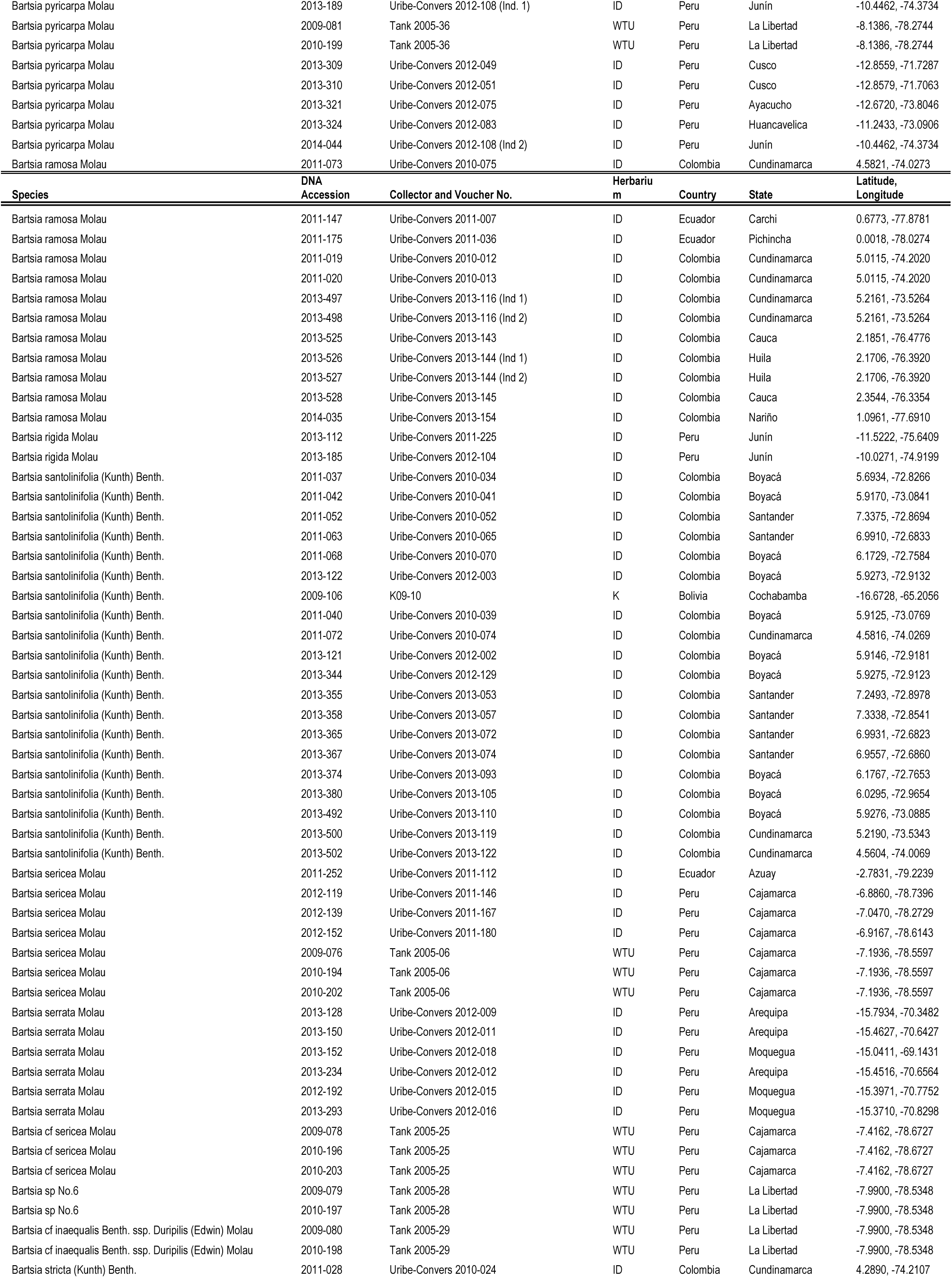
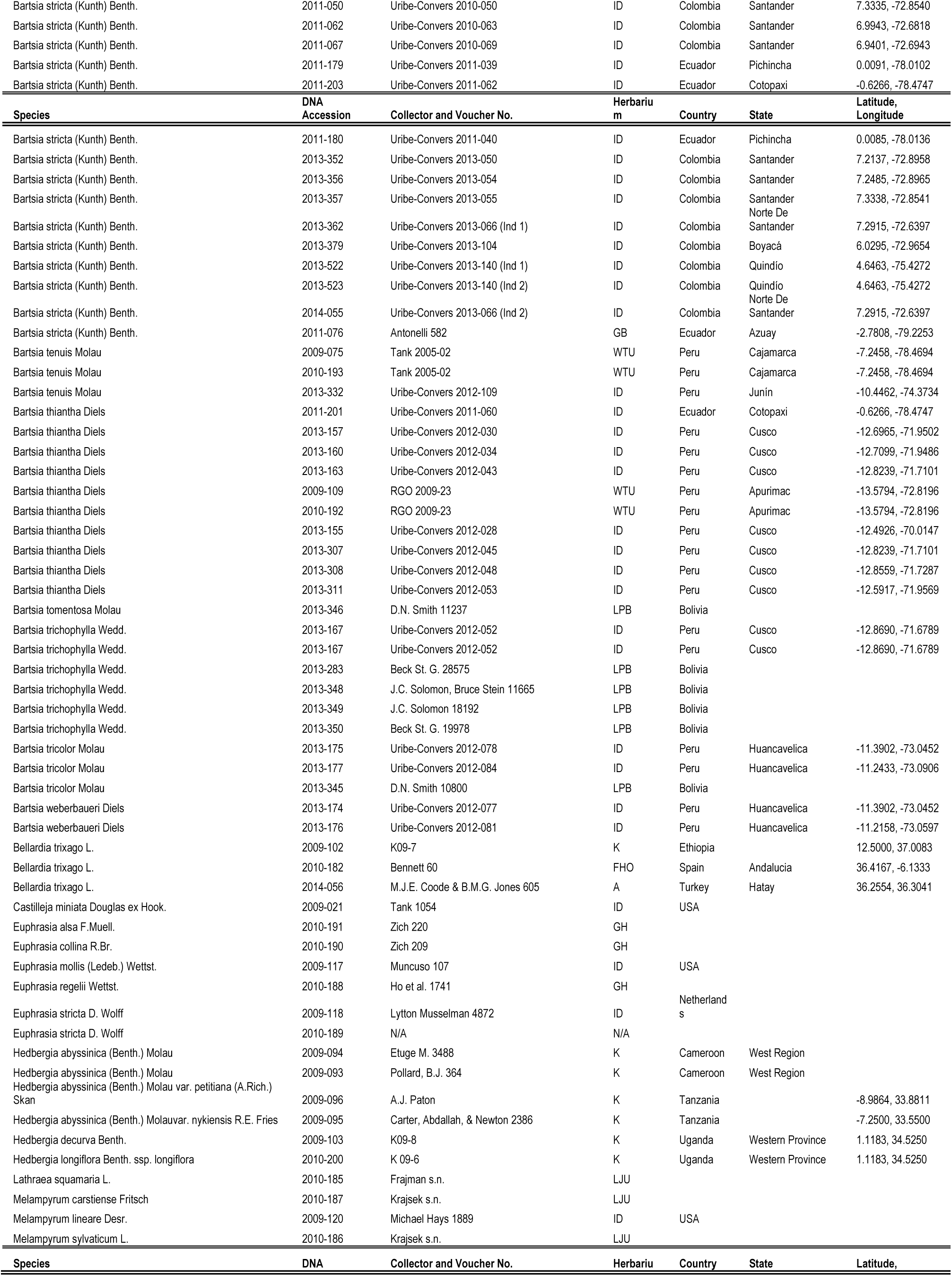
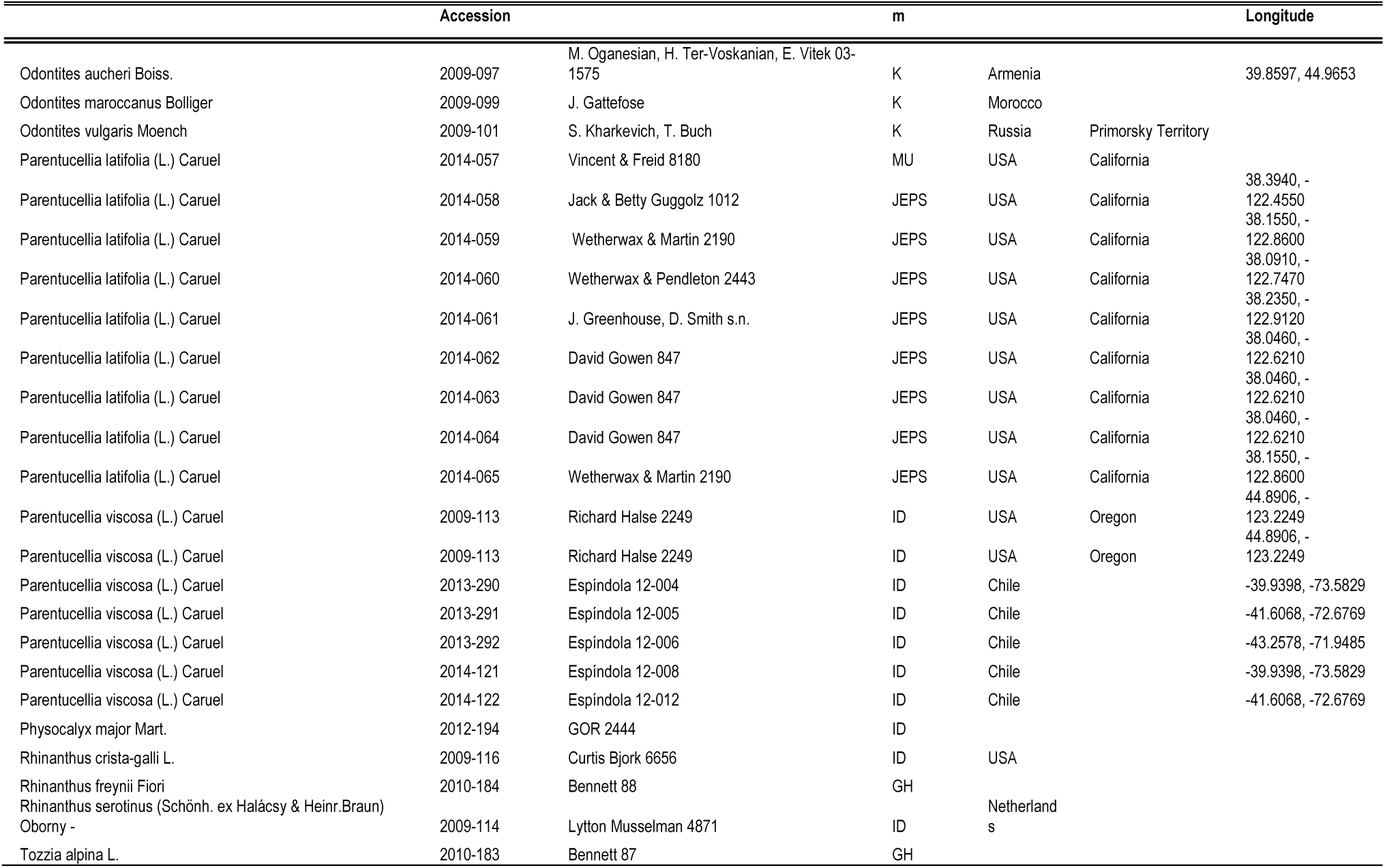
Samples used in this study. Herbarium codes follow the Index Herbariorum.

## S4 Table. Allele occurrences found for each species in this study

Tab 1: Summary of the number of alleles in every sample for each species, and a percentage of how many of these were recovered with more than two alleles. An estimated ploidy level is also given. A reference table of *Bartsia* species and closely related taxa with published chromosome counts from Molau (1990) is also included.

Tab 2: Summary of the number of alleles found in each locus for every sample, and a percentage of how many loci had this specific number of alleles.

Tab 3: Raw data of the number of alleles found in each locus for every sample.

